# High-throughput *in vivo* screening using barcoded mRNA identifies lipid nanoparticles with extrahepatic tropism for cancer immunotherapy

**DOI:** 10.1101/2025.06.09.658602

**Authors:** Alex G. Hamilton, Ajay S. Thatte, Junchao Xu, Zhangyi Luo, Hannah C. Safford, Kelsey L. Swingle, Jenna Muscat-Rivera, Michael Kegel, Xuexiang Han, Ryann A. Joseph, Amanda M. Murray, Hannah C. Geisler, Ricardo C. Whitaker, Lulu Xue, Roman Spektor, Jilian R. Melamed, Drew Weissman, Michael J. Mitchell

## Abstract

Interest continues to grow in the use of mRNA vaccines for cancer immunotherapy. While effective for immunization against infectious diseases, current clinical lipid nanoparticle (LNP) formulations used for mRNA delivery suffer from off-target accumulation, poor immune transfection, and reactogenicity, limiting their application to cancer immunotherapy. Development of new mRNA LNPs is severely bottlenecked by the LNP discovery process, which is historically low-throughput due to reliance on low-plexity measurements. Here, we develop a next-generation high-throughput *in vivo* mRNA LNP screening platform based on barcoded mRNA (b-mRNA). Using this b-mRNA screening platform to simultaneously evaluate 122 LNPs, we identify novel LNP formulations capable of potent hepatic and extrahepatic transfection. We employ novel biochemical characterization techniques to analyze nanoparticle protein corona formation with single-particle resolution and gain insight into the influence of protein adsorption on hepatic and splenic transfection. We evaluate a lead LNP candidate for therapeutic cancer vaccination in a syngeneic mouse model of melanoma and demonstrate a significant reduction in tumor burden and increase in survival compared to a clinical mRNA LNP formulation. Together, our results demonstrate the value of advanced LNP screening and characterization techniques for the development of next-generation mRNA therapeutics and vaccines.

## Introduction

mRNA vaccines have emerged as a revolutionary immunization technology following the immense clinical success of the Pfizer/BioNTech and Moderna COVID-19 vaccines.^1^ The rapid development and approval of these vaccines demonstrated the strengths of mRNA technology, which by nature is substantially more modular than previous subunit vaccines.^2,3^ In addition to improving manufacturability, this modularity enables the ready generalization of mRNA vaccines to other disease targets, work which is well represented by myriad ongoing clinical trials and the recent U.S. Food and Drug Administration (FDA) approval of mRESVIA, an mRNA vaccine against respiratory syncytial virus (RSV).^4–6^ An emerging area of interest is the development of mRNA-based cancer vaccines, especially as the modularity and rapid production capacity of mRNA vaccines is well-suited for the development of personalized neoantigen vaccines.^7^

The promise of mRNA vaccines can only be realized through successful delivery of mRNA to cells, a significant engineering challenge that has taken decades to overcome.^4^ The premier mRNA delivery technology is lipid nanoparticles (LNPs), which possess adjuvant properties that make them particularly appealing for mRNA vaccines.^8^ However, while current mRNA LNP formulations have been hugely successful in the clinic, they are not without their shortcomings. Notable limitations of current clinical mRNA LNPs include nonspecific accumulation — primary in the liver — and poor transfection of immune cells.^9^ As the primary mRNA delivery target for immunotherapy is immune cells in secondary lymphoid organs, these propensities lead to a need to develop new mRNA LNP formulations with improved transfection profiles for cancer immunotherapy.^10^

The typical mRNA LNP development pipeline relies heavily on screening approaches based on reporter genes encoding bioluminescent or fluorescent proteins.^11^ While effective, LNP screening using reporter genes is inherently low-throughput due to limited multiplexability, limiting the speed of the LNP discovery process. To overcome this challenge, our group and others have previously reported on the development of high-throughput screening (HTS) methodologies based on next-generation sequencing (NGS) for the rapid evaluation of large LNP libraries directly *in vivo*.^12,13^ In these methods, each LNP formulation tested is formulated containing a distinct nucleic acid sequence, each of which differs only in a short “barcode” sequence. This polymorphism enables matching of delivered cargo to the carrier that transported it there, and highly multiplexed NGS can be used to evaluate the accumulation profiles of the barcoded payloads and hence the LNPs in question. These HTS approaches can therefore greatly accelerate the LNP discovery process by allowing the simultaneous *in vivo* evaluation of dozens or even hundreds of LNP formulations in a single cohort of animals. High-throughput LNP screening studies typically employ barcoded DNA (b-DNA), either solely encapsulated or co-encapsulated with mRNA to more closely mimic the properties of mRNA LNPs.^11,14,15^ Our group has previously reported the development of an HTS platform based on barcoded mRNA (b-mRNA) in a proof-of-concept study, demonstrating improved screening fidelity compared to b-DNA screening of the same LNPs.^16^ A new generation of b-mRNA screening technology would help accelerate the development of new vaccines and therapeutics for use in cancer immunotherapy and beyond.

While current efforts in the development of novel mRNA LNPs are bottlenecked by the screening process, there is also a growing need to develop principles to direct the design of future nanoparticle libraries. Key to the production of these design principles is a greater understanding of phenomena driving *in vivo* tropism and transfection of current mRNA LNPs. HTS is well-suited to produce large datasets describing these characteristics of LNPs, and our group has previously leveraged HTS approaches to begin addressing these questions.^11,17^ It is generally accepted that intravenously (*i*.*v*.) administered RNA LNPs are prone to formation of an apolipoprotein E (ApoE)-rich protein corona due to preferential adsorption of ApoE from serum.^18–20^ Adsorbed ApoE is thought to interact with low-density lipoprotein receptors (LDLRs) expressed by hepatocytes and other cells in the liver, leading to predominantly hepatic LNP uptake and RNA tranfection.^21^ Recent studies have reported the development of novel mRNA LNPs that demonstrate pronounced adsorption of serum proteins other than ApoE, such as β_2_-glycoprotein 1 (β_2_-GPI) and vitronectin (Vtn).^22^ As HTS and other screening approaches enable the discovery of novel LNPs with interesting extrahepatic transfection properties, the continued and broadened analysis of factors such as protein corona formation may help investigators to identify design principles for future LNP development. Ideally, methods for such analysis should be both accessible and high-throughput to facilitate the rapid development of next-generation therapeutics and vaccines.

In this work, we develop and validate a next-generation HTS platform for nucleic acid drug delivery based on b-mRNA. We then employ this HTS platform to screen a large library of 122 distinct LNPs, identifying promising LNP formulations for *in vivo* mRNA delivery to parenchymal and immune cells. After confirming the potency of several novel LNP formulations for both hepatic and extrahepatic mRNA delivery, we evaluate biochemical influences on *in vivo* LNP tropism on a single-particle scale using small particle lfow cytometric techniques. Finally, we evaluate our lead mRNA LNPs for cancer vaccination in a preclinical melanoma model and demonstrate significantly decreased tumor burden and significantly increased survival time compared to clinical standard mRNA LNP formulations, highlighting the utility of our new b-mRNA screening platform for accelerating mRNA LNP discovery to enable the development of mRNA cancer vaccines.

## Results and Discussion

### Development and validation of a next-generation barcoded mRNA platform

Our group and others have previously reported the use of high-throughput screening approaches based on b-DNA for LNP delivery screening.^11–14^ While effective, these screening approaches suffer from the drawback that LNP physicochemical and delivery properties are strongly influenced by the nature of the encapsulated payload.^23–25^ The issue of cargo-carrier interdependence can be partially circumvented through the dual encapsulation of mRNA with low amounts of b-DNA to produce more “mRNA-like” LNPs, as demonstrated previously.^11,26^ Ultimately, however, b-DNA screening methods track the b-DNA molecule rather than the mRNA cargo of interest. To allow accurate tracking of mRNA cargo distribution in a high-throughput manner, our group has previously reported a proof-of-concept b-mRNA platform.^16^ Using this platform, we demonstrated that moving the barcode sequence to the mRNA molecule was feasible and, moreover, yielded more accurate results than b-DNA screening. In the present work, we sought to establish a next-generation b-mRNA platform for high-fidelity *in vivo* mRNA LNP screening, sufficient for screening large pooled LNP libraries for extrahepatic mRNA delivery.

As error correction is an important consideration for pooled screening approaches, we first employed an *in silico* optimization approach to develop a large pool of 12 nt barcodes with error correction capabilities.^27^ We sorted the resultant list of 1,814 barcodes by edit distance and selected the 200 sequences with the greatest average edit distance (Supplementary Table 1). Based on these optimized sequences, we created 200 distinct double-stranded DNA (dsDNA) templates for *in vitro* transcription (IVT) differing only in the barcode sequence. Mature b-mRNA contained as primary features a m^7^GpppAmG “cap 1” structure, a 5’ untranslated region (UTR), a codon-optimized enhanced green fluorescent protein (EGFP) coding sequence (CDS), a 3’ UTR, a 12-nt barcode sequence, a short “clamp” sequence, and a poly(A) tail (Figure 1a). b-mRNA products demonstrated uniform electrophoretic mobility (Figure 1b), suggesting comparable size and degree of polyadenylation across sequences. Moreover, Sanger sequencing of selected b-mRNAs and subsequent sequence alignment yielded a strong consensus sequence, with strong alignment to desired sequences within the fixed regions and expected polymorphism within the 12-nt barcode region (Figure 1c). To evaluate the ability of b-mRNA to facilitate transgene expression, we treated HeLa cells with mRNA LNPs containing a pool of all 200 candidate b-mRNAs and compared expression of EGFP to cells treated with the same mRNA LNP formulation encapsulating barcode-free EGFP mRNA. As expected, we observed that EGFP expression did not appear to be influenced by the presence of barcode sequences (Figure 1d).

**Figure 1.**
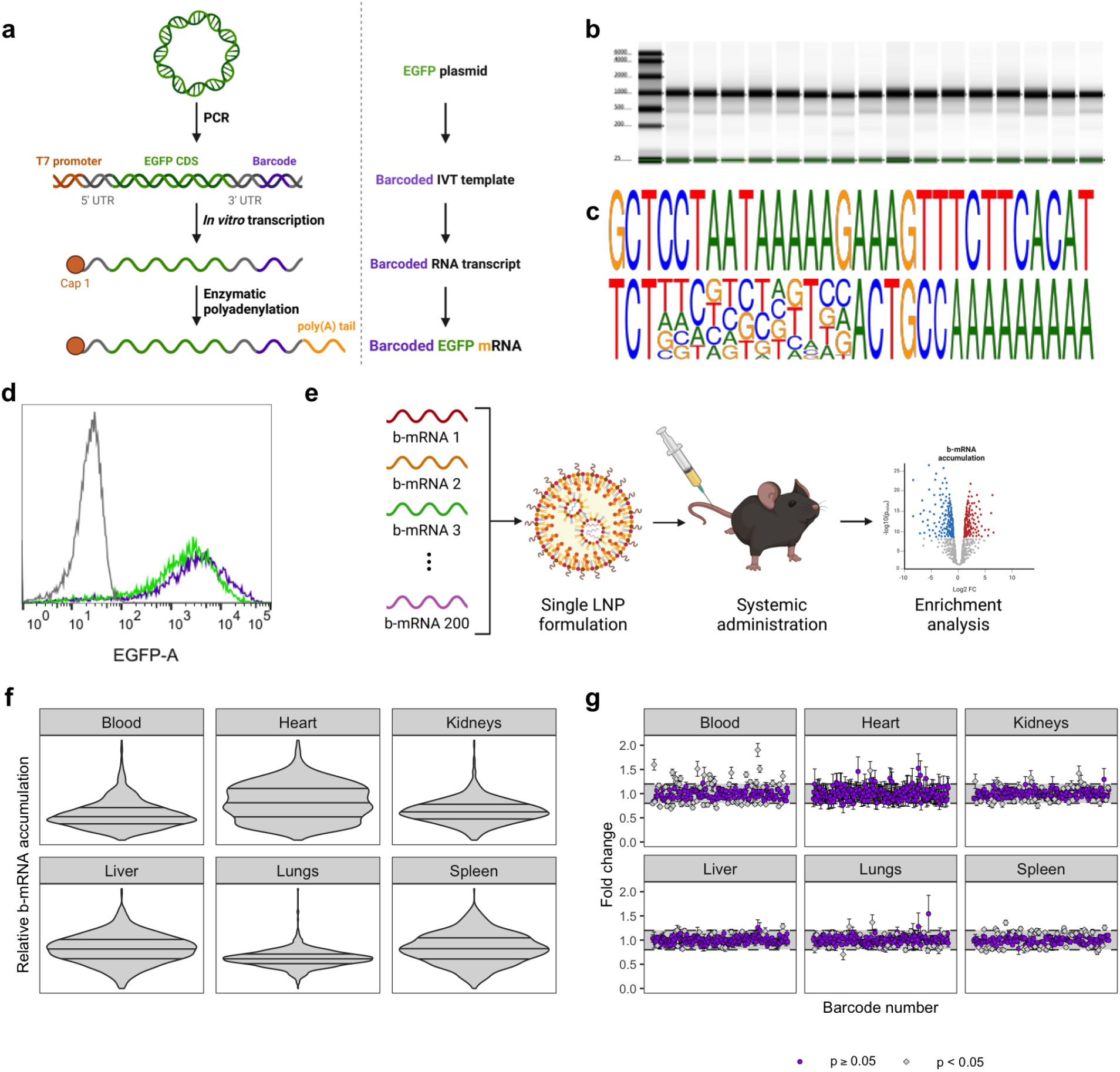
Development of a next-generation barcoded mRNA platform. a. Schematic overview of b-mRNA synthesis through sequential PCR, IVT, and enzymatic polyadenylation. **b**. Representative data from Agilent TapeStation RNA electrophoresis demonstrating integrity and uniform size of b-mRNA products. **c**. Logo visualization of sequence alignment from Sanger sequencing of randomly selected b-mRNAs demonstrating strong alignment outside of the 12 nt polymorphic barcode region. **d**. Representative histogram of EGFP expression demonstrating indistinguishable transgene expression following transfection with equal amounts of EGFP mRNA (green) or EGFP b-mRNA (purple). Data from HeLa cells treated with saline are shown in gray. **e**. Schematic overview of procedure for narrowing of the pool of 200 b-mRNAs to avoid bias from unwanted cellular interactions. **f**. Violin plots illustrating relative homogeneity of b-mRNA accumulation within each tissue of interest. **g**. Enrichment analysis results for selection of b-mRNA candidates from pool narrowing experiments. b-mRNAs demonstrating 20% enrichment or depletion (outside the shaded area) were excluded from later use.

As RNA in the cellular compartment is subject to a plethora of sequence-dependent interactions and regulatory pathways,^28,29^ we next sought to eliminate any b-mRNAs demonstrating differential performance across tissues or cell types. In particular, as endogenous microRNA (miRNA) binding sites are generally considered to be 6 nt to 8 nt in length,^30^ we anticipated the possibility of differential miRNA binding to b-mRNAs within the 12-nt barcode region. As the prevalence of miRNA species differs across tissues and cell types even under homeostatic conditions,^31–33^ differential miRNA binding to b-mRNA could be expected to lead to bias in screening results if overlooked. To avoid screening bias arising from barcode polymorphism, we performed an *in vivo* study of b-mRNA accumulation (Figure 1e). We pooled all 200 b-mRNAs together and encapsulated this pooled library into a single LNP formulation, which we then administered systemically to healthy C57BL/6 mice. We collected peripheral blood, hearts, kidneys, livers, lungs, and spleens from treated mice to assess bioaccumulation of each b-mRNA *via* NGS. As expected, distributions of relative b-mRNA accumulation were largely symmetric and central (Figure 1f). We performed a formal enrichment analysis to identify and eliminate b-mRNAs demonstrating statistically significant enrichment or depletion, suggestive of unwanted interactions with endogenous cellular machinery (Figure 1g), yielding a library of 134 b-mRNAs suitable for pooled *in vivo* mRNA delivery screening (Supplementary Table 2).

### High-throughput *in vivo* screening of LNP-mediated cellular mRNA delivery

Having developed a platform for high-throughput *in vivo* LNP screening, we set out to investigate cellular mRNA delivery facilitated by novel mRNA LNPs. Our group has previously reported that, under systemic administration, ionizable lipid structure plays a pivotal role in cellular tropism.^11^ We therefore opted to screen a large library of novel ionizable lipids, largely focused on a set of biodegradable lipids easily synthesized through a “plug-and-play” strategy from commercial reagents. We have observed strong liver transfection by a subset of these lipids and reasoned that a more detailed investigation of tropism would yield interesting findings in the liver and beyond.^34^ We synthesized 120 “plug-and-play” ionizable lipids and used them to formulate b-mRNA LNPs. We also included several field-standard and industry-standard ionizable lipids as controls, including C12-200,^35^ cKK-E12,^36^ DLin-MC3-DMA,^37^ 306Oi10,^38^ SM-102,^39^ and ALC-0315.^40^ We further included the ionizable lipids previously reported by our group C14-482, C14-488, C16-488, C12-494, C14-494, C16-494, and C14-c494, each of which has shown promise across various RNA delivery applications.^11,41–43^ All told, we synthesized 133 b-mRNA LNPs for evaluation (Supplementary Table 3). We pooled 122 resultant LNPs demonstrating effective mRNA encapsulation (Supplementary Figure 1) with a “naked” b-mRNA, then administered this pool systemically to both C57BL/6 and *APOE*^-/-^ mice to assess ApoE-dependent and -independent mRNA delivery. We collected lungs, spleens, livers, and peripheral blood from treated mice and performed fluorescence-activated cell sorting (FACS) to isolate cell populations of interest before preparing libraries for sequencing *via* NGS.

Using NGS data, we analyzed the biodistribution profile of b-mRNA LNPs within our library. In wild-type mice, several candidate LNP formulations demonstrated both hepatic and extrahepatic tropism (Figure 2a). In *APOE*^-/-^ mice, more standout LNP formulations appeared, likely due to the absence of ApoE-mediated LNP clearance in the liver (Figure 2b).^18^ Across both genotypes, we observed only modest specificity of delivery, with uptake by a given cell type generally correlating well with uptake by other cell types, with the main exceptions of liver endothelium in wild-type mice and lung leukocytes in *APOE*^-/-^ mice (Figure 2c–d). More interestingly, however, we observed generally strong correlation between accumulation profiles in wild-type and knockout mice, suggestive of ApoE-independent hepatic and extrahepatic transfection (Figure 2e). Based on NGS results, we selected LNPs 9, 37, 94, 96, 97, and 112 for further evaluation. These LNPs contained the biodegradable ionizable lipids 2D8, 7D4, 17D8i, 17D9.2, 18D4, and 20D8i, respectively.^34^

**Figure 2.**
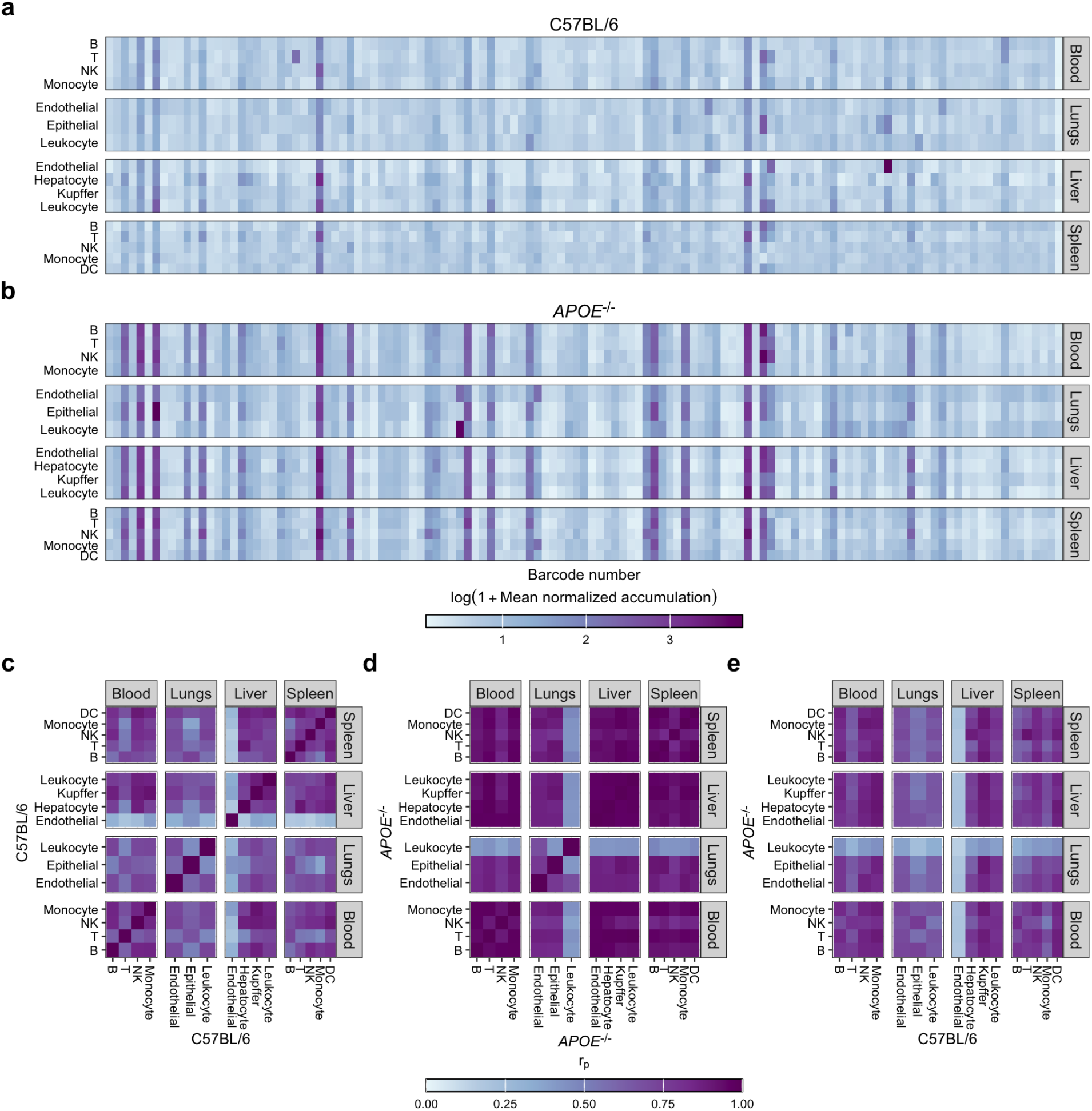
Overview of barcoded mRNA LNP screening results. a–b. Mean normalized intracellular b-mRNA accumulation in cell subsets in the blood, lungs, liver, and spleen of C57BL/6J (**a**) and *APOE*^-/-^ mice. **c–e**. Pairwise Pearson correlation coefficients for b-mRNA accumulation in each sorted cell population in C57BL/6 mice (**c**), *APOE*^-/-^ mice (**d**), and C57BL/6J mice *vs. APOE*^-/-^ mice.

### Validation of lead LNP candidates for extrahepatic mRNA transfection

To confirm transfection of lead mRNA LNP candidates identified by our high-throughput b-mRNA screen, we performed low-throughput validation studies. We first assessed the extrahepatic transfection capabilities of our LNPs at the organ level. We treated C57BL/6 mice with the selected LNP candidates encapsulating NanoLuc mRNA at a low dose and assessed organ transfection using bioluminescence imaging (Figure 3a). Interestingly, all novel LNP candidates tested demonstrated clear splenic transfection, with most demonstrating minimal liver transfection. To aid in evaluating tropism of candidate mRNA LNP, we quantified splenic *vs*. hepatic luminescence signal to classify LNPs as spleen-or liver-tropic (Figure 3b). Interestingly, LNPs 9, 37, 94, 96, and 112 demonstrated splenic transfection with relatively little liver transfection, causing us to classify them as spleen-tropic. LNPs 97 demonstrated strong hepatic transfection, causing us to classify it as liver-tropic. As expected, SM-102 LNPs were also classified as liver-tropic, though they did additionally demonstrate strong splenic transfection. Excitingly, LNP 112 achieved extremely strong splenic transfection, with comparable spleen luminescence to SM-102 LNPs and over an order-of-magnitude decrease in liver luminescence, sufficient to shift its classification to spleen-tropic. As such, LNP 112 emerged from first-pass validation as our lead LNP candidate for *in vivo* reprogramming of splenocytes. Having established the potency of lead mRNA LNP candidates, we next sought to investigate cellular tropism in greater detail. To this end, we treated C57BL/6 mice with fluorescently labeled EGFP mRNA LNPs and performed flow cytometry to analyze accumulation at the cellular level. In the liver, we observed generally high LNP accumulation in Kupffer cells, with lower degrees of accumulation occurring in hepatocytes and non-Kupffer leukocytes (Figure 3c–d). For most LNPs tested, we generally observed greater transfection of Kupffer cells than other liver cell types, which agrees well with observed accumulation data (Figure 3e–f). However, SM-102 LNPs demonstrated very strong transfection of hepatocytes — over 50% on average — and appreciable but less marked transfection of liver endothelial cells. Strikingly, LNP 97 mediated even stronger hepatic transfection than SM-102 LNPs, attaining nearly 80% mean hepatocyte transfection and roughly 20% transfection of liver endothelial cells. These impressive hepatic transfection characteristics make LNP 97 a promising candidate for treating disorders of the liver and may further make it an interesting prospect for treating autoimmune disorders by leveraging the tolerizing environment of the liver.^44^

**Figure 3.**
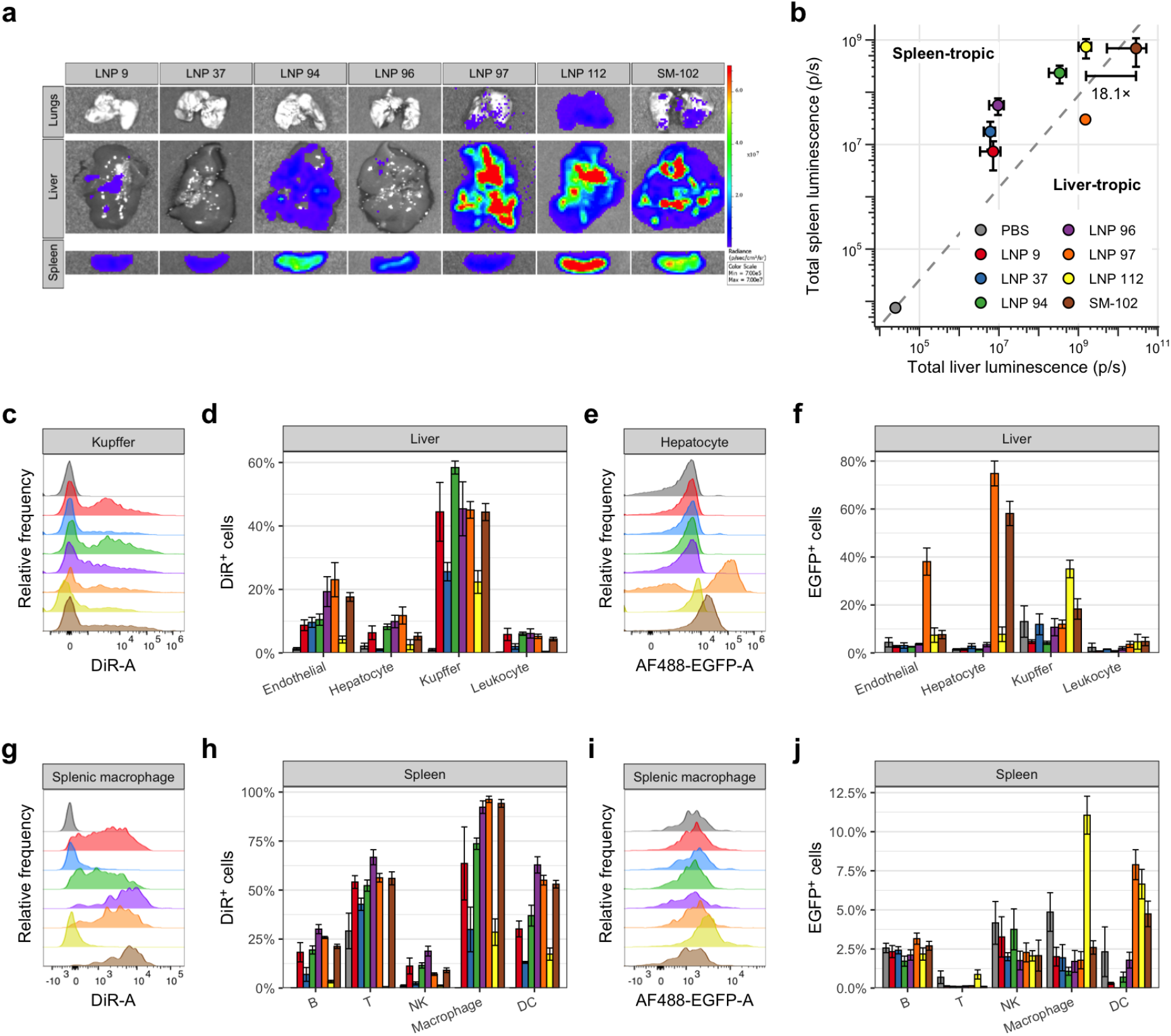
Validation of lead mRNA LNP candidates. a–b. Representative NanoLuc luminescence images (a) and region-of-interest quantification (b) for lead candidate LNPs evaluated. C57BL/6J mice were treated with mRNA LNPs at a dose of 0.1 mg/kg encapsulated mRNA to verify transfection and tropism. **c–d**. Representative flow cytometry histograms of DiR signal in Kupffer cells (c) and detailed quantification of DiR positivity rate in major cell populations of the liver (d) following treatment with DiRlabeled mRNA LNPs. **e–f**. Representative flow cytometry histograms of EGFP expression in Kupffer cells (e) and detailed quantification of EGFP positivity rate in major cell populations of the liver (f) following treatment with lead EGFP mRNA LNPs. **g–h**. Representative flow cytometry histograms of DiR signal in splenic macrophages (g) and detailed quantification of DiR positivity rate in major cell populations of the spleen (h) following treatment with DiR-labeled mRNA LNPs. **i–j**. Representative flow cytometry histograms of EGFP expression in splenic macrophages (i) and detailed quantification of EGFP positivity rate in major cell populations of the spleen (j) following treatment with lead EGFP mRNA LNPs. C57BL/6 mice were treated with DiR-labeled EGFP mRNA LNPs at a dose of 0.6 mg/kg encapsulated mRNA to investigate detailed cellular tropism and transfection.

In the spleen, our tested LNPs largely accumulated in myeloid cells (Figure 3g–h). LNP 96, LNP 97, and SM-102 LNPs demonstrated indistinguishable accumulation in splenic macrophages, each reaching in excess of 70%. LNPs 9 and 94 each accumulated in roughly 40% of splenic macrophages, while LNPs 37 and 112 demonstrated substantially lower accumulation. LNP 96, LNP 97, and SM-102 similarly demonstrated the greatest accumulation in dendritic cells (DCs) of LNPs tested, each reaching at least 30%, while LNPs 9 and 94 again demonstrated lower accumulation and LNPs 37 and 112 again demonstrated lower signal. Strikingly, LNP 112 demonstrated remarkable transfection of splenic myeloid cells, attaining over 10% mean transfection of splenic macrophages and over 6% mean transfection of splenic DCs (Figure 3i–j). As with the liver, despite its low accumulation in myeloid cells, LNP 112 demonstrated strong transfection of these cell types in the spleen, an exciting finding for vaccine applications. Altogether, validation experiments demonstrated the potential of lead LNPs for both hepatic and extrahepatic mRNA delivery, identifying LNP 112 as a lead candidate for the *in vivo* engineering of splenocytes for cancer immunotherapy.

### Single-particle assessment of serum protein adsorption to LNPs

Having evaluated the organ and cellular tropism of our mRNA LNPs, we next sought to investigate possible factors driving LNP tropism. The nanoparticle protein corona has been identified as a major determinant of *in vivo* fate.^45^ LNPs in particular are known to interact with ApoE in circulation, which interacts with LDLRs on cells in the liver to endow much of the classical hepatic tropism of LNPs.^18–21^ Research in recent years has identified other candidate serum proteins that may help to direct LNP tropism to extrahepatic targets such as the spleen and the lungs through their participation in the formation of a nanoparticle protein corona.^22^ One of the serum proteins currently thought to be responsible for the spleen tropism of some LNP formulations is β2-GPI.^22,43^ We therefore sought to evaluate the adsorption of both ApoE and β_2_-GPI to the surface of our lead mRNA LNP formulations to better understand the mechanisms behind their tropism.

Previous studies of LNP protein corona formation have largely relied on classical proteomics approaches based on centrifugation and mass spectrometry (MS).^22^ While effective and useful, these assays are labor-intensive, slow, and rely on the skilled interpretation of complex MS datasets — a difficult feat given the diversity of chemical species found in both LNPs and biological fluids. Additionally, these assays only give insight into bulk properties of an aggregate of LNPs, not single-particle resolution. To gain insight into protein adsorption with single-particle resolution in a rapid manner, we developed a novel experimental approach based on small-particle flow cytometry. Based on previous studies analyzing LNP properties at a single-particle level using custom flow devices, we employed a commercially available flow cytometer designed for the detection of extracellular vesicles (EVs) to analyze protein binding to LNPs. We reasoned that leveraging (i) flow cytometry, a widely used technique in the nucleic acid delivery field, and (ii) a commercially available flow cytometer would help to maximize the accessibility of our experimental technique. We fluorescently dyed NanoLuc reporter mRNA and encapsulated it in LNPs formulated containing a small amount of dyed phospholipid. We then fluorescently dyed recombinant mouse ApoE and β_2_-GPI proteins. After establishing suitable parameters for the detection of single LNPs, we incubated fluorescently dyed NanoLuc LNPs with dyed proteins of interest at a fixed molar ratio. After incubation, we diluted samples to minimize further interactions and acquired data using flow cytometry. Adsorption of protein to candidate LNPs resulted in an increase in protein fluorescence, which we used to quantify the proportion of LNPs with adsorbed protein (Figure 4a).

**Figure 4.**
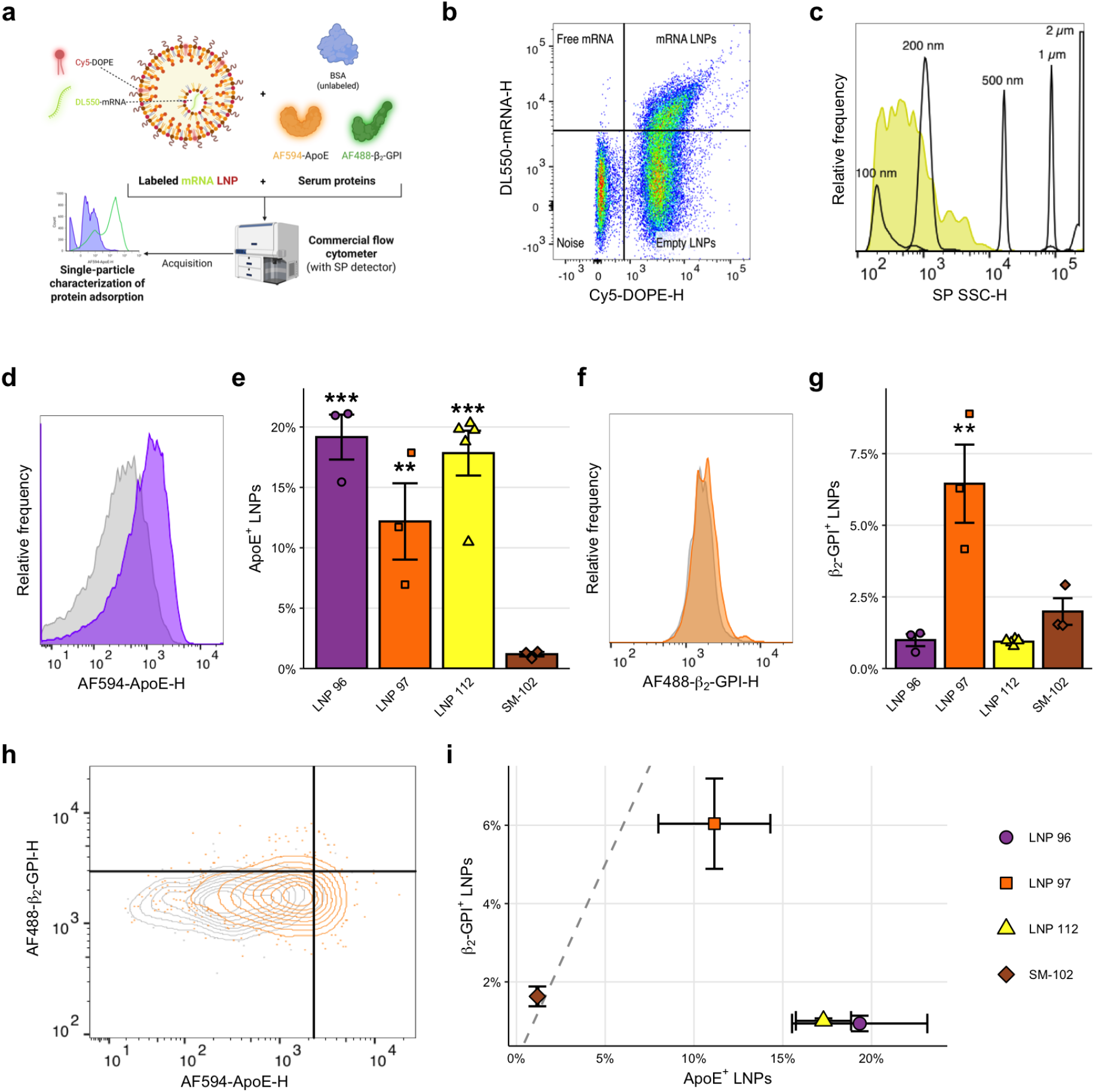
Single-particle assessment of serum protein adsorption to LNPs. a. Schematic overview of experimental approach. Fluorescently labeled LNPs were prepared through the incorporation of Cy5-labeled DOPE and DL550-labeled mRNA. Mouse ApoE and β_2_-GPI proteins were fluorescently labeled with different fluorochromes. Labeled LNPs were then incubated with mouse BSA alone or in combination with fluorescently labeled ApoE and/or β_2_-GPI. A commercial flow cytometer with a small particle detector was then used to analyze samples. **b**. Representative flow cytometry plot of fluorescently labeled LNPs. Dyed lipid and RNA components distinguish free mRNA, empty LNPs, and mRNA-loaded LNPs. **c**. Representative flow cytometry histogram visualizing estimated size of studied particles. Calibration beads (diameters indicated) were used to estimate the approximate size of the LNPs under study. **d–e**. Representative histogram (d) and detailed quantification (e) of ApoE adsorption to single LNPs for mRNA LNP candidates of interest. **f–g**. Representative histogram (f) and detailed quantification (g) of β_2_-GPI adsorption to single LNPs for mRNA LNP candidates of interest. **h**. Representative contour plot visualizing the effects of equimolar incubation of LNPs with fluorescently labeled ApoE and β_2_-GPI (competitive binding). **i**. Detailed quantification of competitive β_2_-GPI adsorption *vs*. ApoE adsorption to single LNPs. Bar and scatter plots represent data from *n* ≥ 3 independent experiments for each LNP formulation. Gray contours and histograms represent LNPs incubated with BSA only. Significance annotations indicate comparisons to SM-102 LNPs. **: *p* < 0.01. ***: *p* < 0.001.

Upon analysis of flow cytometry data, we were able to clearly resolve three distinct event populations: free mRNA, mRNA-loaded LNPs, and empty LNPs (Figure 4b). Using fluorescent polystyrene size calibration beads, we confirmed that the diameter of analyzed particles was generally between 100 nm and 200 nm, in line with dynamic light scattering (DLS) measurements of LNP size (Figure 4c). Having confirmed the validity of our measurements, we then analyzed particle fluorescence following incubation either solely with unlabeled bovine serum albumin (BSA) or with a mixture of unlabeled BSA and fluorescently labeled ApoE or β_2_-GPI. As expected, we generally observed strong adsorption of ApoE to the tested LNP formulations (Figure 4d–e). Somewhat surprisingly, however, SM-102 LNPs demonstrated minimal adsorption of ApoE. Our group has previously reported that ApoE readily binds to 1,2-distearoyl-*sn*-glycero-3-phosphoethanolamine (DOPE)-containing LNPs, which may help explain this finding.^17^ Importantly, as some leukocytes express LDLRs, we do not expect adsorption of ApoE to contribute solely to hepatic tropism: instead, it is likely that increased adsorption of ApoE enhances trafficking to both the spleen and the liver, to possibly differential extents.^46,47^

The mechanism(s) by which β_2_-GPI adsorption leads to spleen tropism, if any, remain unclear, though this effect has been postulated to stem from the presence of externalized anionic phospholipids in the spleen.^22^ However, our group has previously shown that increased β_2_-GPI binding capacity may improve transfection of splenocytes, with more modest increases in hepatic transfection.^43^ We therefore anticipated increased binding affinity for β_2_-GPI for our spleen-tropic LNP formulations. Interestingly, however, we observed the strongest β_2_-GPI adsorption for LNP 97, our strongest liver-tropic LNP, which demonstrated over 5-fold greater β_2_-GPI adsorption than any of our spleen-tropic LNP formulations (Figure 4f–g).

To validate our findings from single-protein experiments, we next performed competitive binding experiments wherein we incubated dyed LNPs with equimolar amounts of dyed ApoE and β_2_-GPI and assessed the adsorption of both proteins to the LNP surface. Competitive binding results generally recapitulated those from single-protein investigations, though we observed an expected decrease in protein binding in some cases (Figure 4h–i). In all, our results suggest a nuanced relationship between serum protein adsorption and transfection of hepatic and extrahepatic targets and support previous findings of differential adsorption of serum proteins based on ionizable lipid structure.^48^

### Evaluation of mRNA LNP therapeutic effect in a syngeneic mouse model of melanoma

Based on the strong performance of LNP 112 for transfection of splenic antigen-presenting cells (APCs), we sought to evaluate its potential for use in a cancer vaccine application. To this end, we established a syngeneic mouse melanoma model by inoculating C57BL/6 mice of mixed sex with B16-OVA melanoma cells. After allowing tumors to engraft, we treated mice with one of four treatments at a 5 d prime-boost interval (Figure 5a): (i) phosphate-buffered saline (PBS) (negative control), (ii) LNP 112 encapsulating antigen-irrelevant firefly luciferase (FLuc) mRNA (vehicle control), (iii) LNP 112 encapsulating ovalbumin (OVA) antigen mRNA, or (iv) SM-102 LNPs encapsulating OVA mRNA (clinical control). We closely monitored mouse body weight (Supplementary Figure 5), tumor volume, and survival over the course of 60 d. Excitingly, LNP 112 encapsulating OVA mRNA demonstrated a strong therapeutic effect, substantially reducing tumor burden compared to any other treatment group over the course of 21 d (Figure 5b) and significantly improving survival (Figure 5c), with a median survival time of 31 d. Notably, of the 12 mice treated with LNP 112 containing OVA mRNA, 2 mice entirely cleared their tumors following the boost dose.

**Figure 5.**
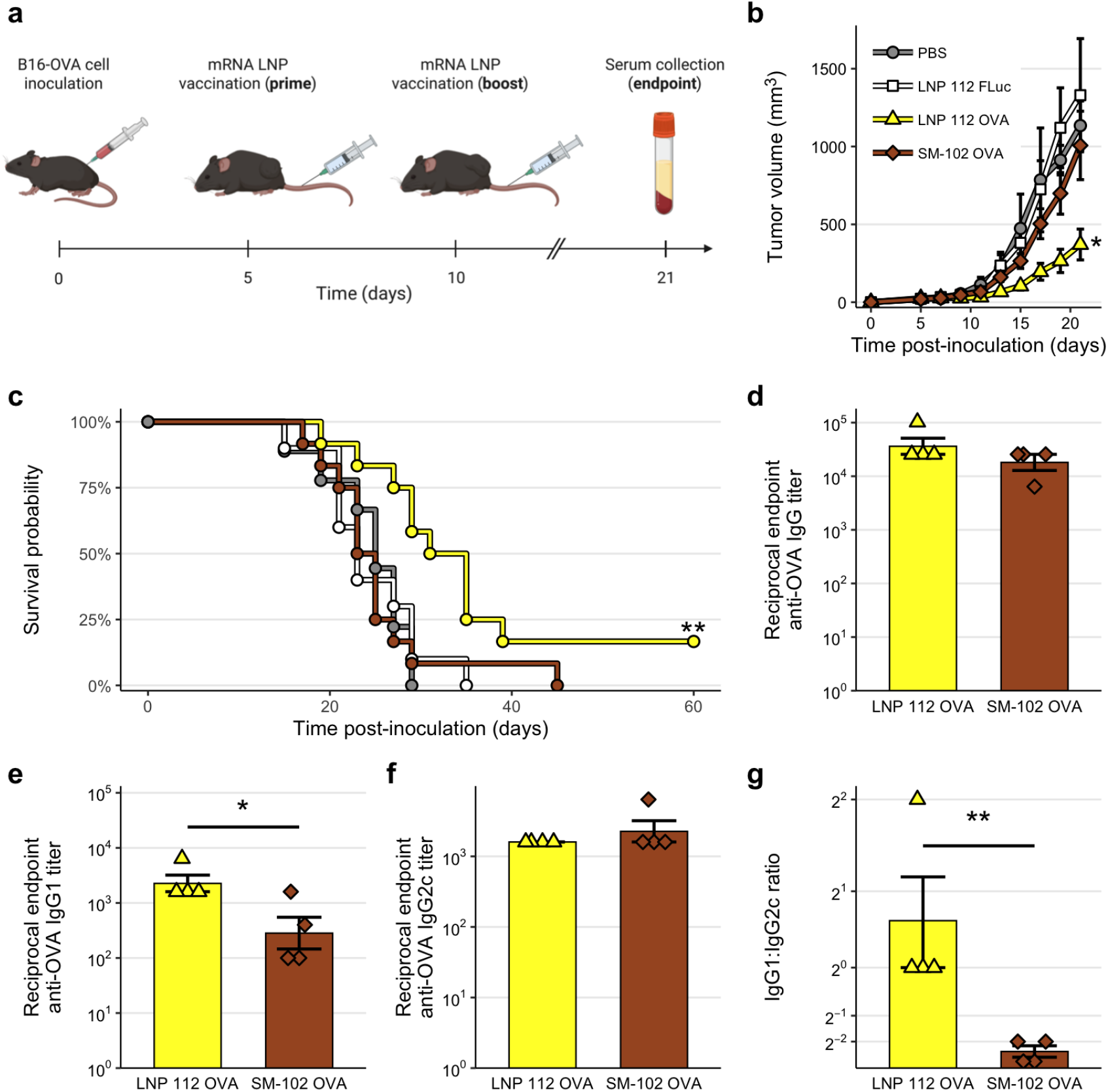
Evaluation of lead mRNA LNP candidate in a preclinical therapeutic cancer vaccine model. a–b. Tumor burden (a) and survival curves (b) from a murine syngeneic melanoma solid tumor disease model. Tumors were established by inoculating C57BL/6 mice with B16-OVA cells and mice were subsequently treated *i*.*v*. with PBS, LNP 112 encapsulating FLuc mRNA, LNP 112 encapsulating OVA mRNA, or SM-102 LNPs encapsulating OVA mRNA with a 5 d prime-boost interval. **c–e**. Reciprocal endpoint total anti-OVA IgG (c), anti-OVA IgG1 (d), and anti-OVA IgG2c (e) titers of mice treated with OVA mRNA LNPs. **f**. Ratio of endpoint reciprocal anti-OVA IgG1 to anti-OVA IgG2c titer in sera of mice treated with OVA mRNA LNPs. Tumor burden and survival data are summarized from *n* ≥ 9 mice per treatment group, while antibody titer data represent *n* = 4 animals per group. Significance annotations without bars indicate comparisons to PBS-treated mice. *: *p* < 0.05. **: *p* < 0.01.

To characterize the immune response to vaccination with LNP 112, we collected mouse sera and splenocytes 7 d following the boost dose. To characterize the humoral immune response, we quantified reciprocal endpoint anti-OVA antibody titers using enzyme-linked immunosorbent assays (ELISAs). Interestingly, despite the clearly superior performance of LNP 112 to SM-102 LNPs, total anti-OVA immunoglobulin G (IgG) appeared to be comparable between the groups (Figure 5d). However, analysis of antibody isotypes revealed significantly higher endpoint antigen-specific IgG1 titers for the LNP 112-treated group (Figure 5e), with comparable endpoint IgG2c titers (Figure 5f). As a result, quantified endpoint antigen-specific IgG1:IgG2c titer ratios were significantly greater for mice treated with LNP 112 compared to those treated with SM-102 LNPs (Figure 5g). As IgG1 production is indicative of a type 2 helper T (Th2) immune response while IgG2c production indicates a type 1 helper T (Th1) response, these results suggest a greater Th2 bias in mice vaccinated with LNP 112 compared to SM-102 LNPs. The improved efficacy of LNP 112 in evoking anti-tumor responses may therefore be due to enhanced induction of humoral immunity compared to SM-102 LNPs.

## Conclusions

In this work, we report the development of a next-generation b-mRNA screening platform with strong error correction capabilities for high-fidelity mRNA delivery screening. We formulated a library of 133 distinct mRNA LNPs, each containing a distinct ionizable lipid. From screening in both wild-type and *APOE*^-/-^ mice, we identified several promising LNPs for both hepatic and extrahepatic mRNA transfection, each based on highly accessible “plug-and-play” ionizable lipids. We employed low-throughput counterscreening experiments to validate functional mRNA transfection on both the tissue and cellular levels, identifying LNP 96 as an exceptionally strong hepatic transfection candidate and LNP 112 as a promising candidate for splenic myeloid cell reprogramming. To investigate potential influences of the nanoparticle protein corona on LNP tropism, we developed a novel experimental approach for single-particle analysis of protein adsorption. We employed this technique to analyze the adsorption of ApoE and β_2_-GPI, serum proteins generally regarded as pivotal in dictating *in vivo* LNP fate, to our lead LNP formulations, demonstrating differential adsorption, with high ApoE adsorption to top spleen-tropic formulations and strong β_2_-GPI adsorption to our most liver-tropic formulation. In a preclinical melanoma model, we demonstrated that the spleen-tropic LNP 112 was effective in reducing disease burden and prolonging survival compared to LNPs formulated with a clinical ionizable lipid. Analysis of the humoral immune response suggested a greater induction of humoral immunity following cancer vaccination compared to clinical LNPs. In all, our data demonstrate the value of b-mRNA as an *in vivo* HTS platform for rapid discovery of mRNA LNPs for use in therapeutics and vaccines. We anticipate the adaptation of this screening approach for the evaluation of large LNP libraries for a variety of extrahepatic protein replacement, vaccine, gene editing, and immunotherapy applications. Moreover, the combination of our HTS data, validation experiments, and single-particle analysis of ApoE and β_2_-GPI adsorption paints a nuanced picture of protein corona formation and *in vivo* interactions and highlights a need for more widespread characterization of protein adsorption to advance the field’s understanding of LNP fate, work which we hope our reported analytical techniques will further help to accelerate.

## Methods

### High-throughput screening platform development

#### Barcode sequence design

Barcode sequences were designed using *in silico* optimization essentially as described previously.^27^ The Conway lexicographic algorithm was used to generate the initial set of barcode sequences, which was refined to yield a pool of 12-nt barcodes with a minimum “sequence Levenshtein” distance of 4.^27^ The resultant set of barcodes guarantees the detection of up to three insertions, deletions, and substitutions in a DNA-anchored barcode context and the correction of at least one sequencing error.

#### Barcoded mRNA synthesis

Barcoded EGFP mRNA was derived from an IVT plasmid containing a 5’ UTR derived from tobacco etch virus (TEV), a codon-optimized EGFP CDS, and a 3’ UTR derived from *Xenopus laevis* beta globin. To produce barcoded template, a constant forward primer and distinct reverse primers were used in the polymerase chain reaction (PCR) to introduce a CleanCap AG-compatible T7 promoter sequence 5’ of the TEV UTR and to add a 12-nt barcode sequence and short spacer sequence 3’ of the globin UTR. Solid-phase reversible immobilization (SPRI) beads were used to purify barcoded dsDNA templates for IVT. b-mRNA precursor was synthesized using IVT with co-transcriptional capping using the CleanCap AG trinucleotide cap 1 analog. Uridine residues were fully substituted with N^1^-methylpseudouridine (m^1^Ψ). Following SPRI purification, IVT products were enzymatically polyadenylated using *E. coli* poly(A) polymerase. Mature b-mRNA was purified using SPRI beads and RNA concentration was quantified by absorbance measurements at a wavelength of 260 nm. b-mRNA was diluted to 1 mg/mL and stored at −80 °C for later use.

### LNP synthesis and characterization

LNPs were produced *via* microfluidic mixing essentially as described previously.^49^ Briefly, all lipid components were combined in ethanol, mRNA was placed in acidic citrate buffer, and the phases were combined using chaotic mixing in a microfluidic device containing staggered herringbone micromixers at an aqueous:organic flow rate ratio of 3:1. Resultant LNPs were dialyzed against PBS for 2 h and stored at 4 °C for later use.

mRNA entrapment was measured using a RiboGreen assay essentially as described previously.^25^ Briefly, LNPs were diluted 100-fold in tris-EDTA (TE) buffer or TE buffer containing 0.1% Triton X-100. RNA standards of known concentration were plated along with diluted LNP samples in a well plate and fluorescence detection was used to determine RNA concentration in each sample. Encapsulated mRNA concentration was determined by subtracting the concentration in TE buffer (free mRNA) from that in buffer containing detergent (total mRNA). mRNA entrapment efficiency was determined as the ratio of encapsulated RNA concentration to total RNA concentration.

LNP size distributions were assessed *via* DLS using a DynaPro plate reader (Wyatt Technology Corporation, Santa Barbara, CA) following 10-fold dilution in PBS.

### Flow cytometry and FACS

#### Primary cell isolation

For analysis of circulating immune cells, peripheral blood was collected into microcentrifuge tubes precoated with EDTA to prevent clotting. Mice were thoroughly perfused with PBS containing EDTA before dissection, and lungs, livers, and spleens were removed and placed in Hank’s buffered salt solution (HBSS) containing collagenase IV and DNase I for digestion. After enzymatic digestion, organs were mechanically digested through 100 µm strainers. All samples containing red blood cells were treated with ammonium-chloride-potassium (ACK) lysing buffer. Processed liver samples were centrifuged through a discontinuous Percoll gradient to remove debris.

#### Cell staining

All antibodies used for fluorescence detection were obtained from BioLegend. A list of clones used is provided in the Supplementary Information (Supplementary Table 4). Cells were blocked using anti-CD16/CD32 antibodies and stained using the Zombie UV amine-reactive dye following the manufacturer’s instructions. After quenching with PBS containing BSA, cells were stained with extracellular antibodies for 30 min. For studies requiring intracellular staining, cells were fixed and permeabilized using a Cyto-Fast Fix-Perm Buffer Set according to the manufacturer’s instructions. Cells were then stained with antibodies in permeabilization buffer for 30 min before rinsing and resuspension in PBS.

#### Flow cytometry data acquisition

For cellular flow cytometry experiments, a FACSymphony A3 equipped with UV, violet, blue, yellow-green, and red lasers was employed. In all cases, at least 10,000 events were acquired. Data for compensation were acquired using single-stained controls. Fluorescence-minus-one (FMO) controls were employed as needed to assist in data analysis. Flow cytometry data were analyzed using FlowJo version 10.

#### FACS

For b-mRNA LNP screening, FACS was performed to isolate parenchymal and immune cells from processed tissues. A FACSymphony S6 equipped with UV, violet, blue, yellow-green, and red lasers was employed to sort B cells, T cells, NK cells, and monocytes from the blood (Supplementary Figure 2); endothelial cells, epithelial cells, and leukocytes from the lungs (Supplementary Figure 3); hepatocytes, endothelial cells, Kupffer cells, and other leukocytes from the liver (Supplementary Figure 4); and B cells, T cells, NK cells, monocytes, and DCs from the spleen.

### NGS library preparation and sequencing

#### RNA isolation

Total RNA was isolated using phenol-chloroform precipitation with Trizol reagent according to the manufacturer’s instructions. Tissue samples were mechanically homogenized in Trizol reagent with a bead mill before processing, while cells were sorted directly into Trizol LS reagent. For samples with low RNA content (*e*.*g*., sorted cells, LNPs), RNA was co-precipitated using 20 µg of glycogen as a carrier. RNA pellets were resuspended in TE buffer with low EDTA content for downstream use.

#### First-strand cDNA synthesis

First-strand complementary DNA (cDNA) synthesis was performed using the ProtoScript II system according to the manufacturer’s instructions. Reverse transcription (RT) oligos were based on the common oligo(dT) method for RT of eukaryotic mRNA and additionally featured (i) a 3’ “clamp” complementary to the barcode-tail spacer region, (ii) a 5’ 12-nt unique molecular identifier (UMI), and (iii) the Illumina Nextera Read 2 sequence.

#### cDNA amplification and index/adapter ligation

To produce final NGS libraries, nested PCR was employed. cDNA was first amplified using a forward primer against either the 3’ portion of the EGFP CDS (for b-mRNA pool narrowing) or the end of the 3’ globin UTR and a reverse primer against the Nextera Read 2 sequence. The forward primer used in this first PCR included an overhanging portion containing the Nextera Read 1 sequence. Nine cycles of PCR were performed to add the Read 1 sequence. Sixteen additional cycles of PCR were performed to add indices (i5/i7) and sequencing adapters (P5/P7) using polymorphic primers against Read 1/Read 2 sequences. NGS libaries were purified using SPRI beads and eluted in TE buffer with low EDTA content for sequencing. Library concentrations were quantified using a Qubit 1× dsDNA assay and DNA was stored at −20 °C for later use.

#### Next-generation sequencing

A NGS library pool was prepared by combining equal masses of each NGS library. The concentration of this library pool was quantified using a Qubit 1× dsDNA assay. The library pool was subjected to quality control using an Agilent BioAnalyzer and sequenced using dual-indexed sequencing on an Illumina MiSeq (b-mRNA pool narrowing) or an Illumina NextSeq 2000 with 10% ΦX174 sequencing control spike-in.

### NGS data analysis and visualization

NGS data were analyzed and visualized similarly to previously described using the R statistical programming language with a number of packages from the Comprehensive R Archive Network (CRAN) and Bioconductor.^11,27,50–62^ Briefly, reads were demultiplexed and trimmed of adapter sequences. Barcode and UMI sequences were then extracted from reads using the UMI-tools Python package.^63^ Barcode sequences were matched to candidate sequences using the DNABarcodes R package,^27^ and UMIs were collapsed to yield read counts for each barcode. Normalized accumulation was calculated as the ratio of output read fraction to input read fraction.^13^ Specifically, read fractions were first computed by dividing the read count for each barcode by the total number of barcode counts within each sample. Each sample read fraction (output) was then divided by the corresponding mean read fraction from the uninjected pool samples (input) to yield normalized accumulation.

For enrichment analysis, Wilcoxon rank-sum tests were performed to compare normalized accumulation for each barcode to the normalized accumulation of all other barcodes using the normal approximation with continuity correction. False discovery rate was controlled using the method of Benjamini and Hochberg.^64^

### Evaluation of lead mRNA LNPs

#### Reporter mRNA synthesis

We employed the NanoLuc engineered luciferase as both a model mRNA and a reporter for validation and mechanistic experiments. EGFP was used as a reporter for studies of cellular transfection charateristics. gBlock dsDNA templates were synthesized containing codon-optimized NanoLuc or EGFP CDSs and the same UTRs as used for b-mRNA production. PCR with overhanging primers was employed to produce dsDNA template for IVT containing a CleanCap AG-compatible T7 promoter sequence and a 100 nt poly(A) tail, which was purified using SPRI beads. Reporter mRNA was synthesized using IVT with co-transcriptional capping with the CleanCap AG trinucleotide cap 1 analog. Uridine residues were fully substituted with m^1^Ψ when using the mRNA as a reporter gene. 1 M urea was used in the IVT reaction as a chaotropic agent to reduce double-stranded RNA (dsRNA) formation.^65^ mRNA was purified using SPRI beads and RNA concentration was quantified by absorbance measurements at a wavelength of 260 nm prior to storage at −20 °C.

#### mRNA LNP transfection validation experiments

To confirm the *in vivo* transfection ability of candidate mRNA LNPs identified by NGS, we performed single-plex validation experiments using NanoLuc. C57BL/6 mice were injected with NanoLuc mRNA LNPs at an encapsulated mRNA dose of 0.1 mg/kg. 6 h later, 220 nmol of fluorofurimazine (FFz) in PBS was administered intraperitoneally (*i*.*p*.) and major mouse organs were dissected and imaged using an *in vivo* imaging system (IVIS) to assess *in vivo* mRNA transfection (PerkinElmer, Shelton, CT).

For single-cell analysis of *in vivo* mRNA LNP transfection and accumulation, we performed flow cytometry experiments. Lead mRNA LNP candidates were reformulated encapsulating EGFP mRNA and fluorescently dyed using lipophilic 1,1’-dioctadecyl-3,3,3’,3’-tetramethylindotricarbocyanine iodide (DiR) at a concentration of 10 µM. C57BL/6 mice were injected with DiR-labeled EGFP mRNA LNPs at an encapsulated mRNA dose of 0.6 mg/kg. 12 h later, mice were sacrificed and livers and spleens were collected for flow cytometric analysis of LNP accumulation and mRNA transfection. To improve EGFP signal strength, an Alexa Fluor 488 (AF488)-conjugated anti-EGFP antibody was employed for intracellular staining.

### Flow cytometric analysis of protein adsorption to LNPs

#### Labeled mRNA synthesis

mRNA was fluorescently tagged using a two-step procedure based on reaction between aminoallyl-modified RNA and amine-reactive *N* -hydroxysuccinimide (NHS) ester-fluorophore conjugates. Aminoallyl-modified mRNA coding for NanoLuc was first synthesized using IVT essentially as described above except using canonical uridine instead of m^1^Ψ. Aminoallyl-UTP (aaUTP) was also incorporated into the nucleoside triphosphate (NTP) mixture at a ratio of 1:1 UTP:aaUTP to produce partially-substituted mRNA. After purification, aminoallyl-modified mRNA was reacted with DyLight 550 (DL550) NHS ester conjugate in basic conditions at an mRNA:fluorophore ratio of 1:200 to produce fluorescently labeled mRNA. Labeled mRNA was purified using SPRI beads, quantified using absorbance measurements, and stored at −20 °C for later use.

#### Fluorescence labeling of serum proteins

Recombinant mouse ApoE and β_2_-GPI proteins were differentially labeled using Alexa Fluor 594 (AF594) and AF488 protein labeling kits, respectively, according to the manufacturer’s instructions. Labeled protein was purified using spin filters according to the manufacturer’s instructions and eluted in PBS. Following purification, protein concentration was quantified using adjusted absorbance measurements before storage at 4 °C.

#### Flow cytometric analysis of single LNPs

LNPs of interest identified by high-throughput screening were reformulated containing fluorescently labeled NanoLuc mRNA and with 0.5% 1,2-distearoyl-*sn*-glycero-3-phosphoethanolamine-*N* -cyanine 5 (Cy5-DOPE). LNP concentration was quantified using static light scattering (SLS). LNPs were diluted to a concentration of 50 pM before incubation, while proteins were diluted to 25 nM. All species underwent a 100-fold dilution before acquisition. Data were acquired on a BD FACSymphony A1 equipped with a small particle detector following MIFlowCyt-EV recommendations for small particle flow cytometry.^66^

### Therapeutic melanoma vaccine model

#### Tumor engraftment and monitoring

C57BL/6 mice of mixed sex were obtained from the Jackson Laboratory. Mice were subcutaneously inoculated on the right flank with 7.5 × 10^5^ B16-OVA mouse melanoma cells. Tumor dimensions were regularly measured using electronic calipers and tumor volume was calculated as 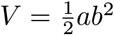, where *a* and *b* represent the largest and smallest tumor dimensions, respectively. Tumor volume and body weight were monitored for up to 60 days.

#### Immunization and evaluation of humoral response

Following successful tumor engraftment, mice were immunized followed a prime-boost strategy at a 5 d interval. Mice were injected *i*.*v*. with mRNA LNPs encapsulating either OVA mRNA or FLuc mRNA at an encapsulated mRNA dose of 0.8 mg/kg. 7 d after the boost dose, mouse sera were collected for analysis.

Reciprocal endpoint antibody titers were measured essentially as described previously.^67^ Briefly, treated 96-well clear polystyrene microplates were coated overnight with a solution of 1 µg/mL OVA protein in PBS. After coating, wells were rinsed with 0.05% Tween 20 in PBS (PBST) and blocked for 1 h using IgG-depleted BSA in PBS. After blocking, wells were again washed with PBST, and diluted endpoint antisera in blocking buffer were plated and allowed to incubate for 2 h. After further rinsing with PBST, diluted horseradish peroxidase (HRP)-conjugated anti-isotype antibody in blocking buffer was added. After rinsing, detection was performed by adding 3,3’,5,5’-tetramethylbenzidine (TMB) substrate for 15 min before quenching with 2 N sulfuric acid. Absorbance was measured using a plate reader at a detection wavelength of 450 nm with a reference wavelength of 650 nm to account for optical effects. Absorbance cutoffs for reciprocal titer determination were determined using the Frey method.^68^

#### Data analysis and statistical inference

All data were analyzed using the R statistical programming language with several packages from CRAN.^50–62,69–74^ The Nix package manager (with pinned Nixpkgs revision a71e967e) was used for all data and dependencies to maximize reproducibility.^75^ Unless otherwise noted, statistical inference was performed on mean responses using one-way analysis of variance (ANOVA) with *post hoc t* tests using the Bonferroni-Holm correction for multiple comparisons. For comparisons involving only two groups, simple unpaired *t* tests were used for inference. For melanoma model survival analysis, pairwise long-rank tests were employed to compare survival curves to PBS-treated mice, and the Bonferroni-Holm correction was applied.^76^

## References

1. Kon, E., Elia, U. & Peer, D. Principles for designing an optimal mRNA lipid nanoparticle vaccine. Current Opinion in Biotechnology 73, 329–336. DOI: 10.1016/j.copbio.2021.09.016 (Feb. 2022).

2. Pardi, N., Hogan, M. J., Porter, F. W. & Weissman, D. mRNA vaccines — a new era in vaccinology. Nature Reviews Drug Discovery 17, 261–279. DOI: 10.1038/nrd.2017.243 (Apr. 2018).

3. Kis, Z., Kontoravdi, C., Dey, A. K., Shattock, R. & Shah, N. Rapid development and deployment of high-volume vaccines for pandemic response. Journal of Advanced Manufacturing and Processing 2, e10060. DOI: 10.1002/amp2.10060 (2020).

4. Cullis, P. R. & Felgner, P. L. The 60-year evolution of lipid nanoparticles for nucleic acid delivery. Nature Reviews Drug Discovery 23, 709–722. DOI: 10.1038/s41573-024-00977-6 (Sept. 2024).

5. Wilson, E., Goswami, J., Baqui, A. H., Doreski, P. A., Perez-Marc, G., Zaman, K., Monroy, J., Duncan, C. J. A., Ujiie, M., Rämet, M., Pérez-Breva, L., Falsey, A. R., Walsh, E. E., Dhar, R., Wilson, L., Du, J., Ghaswalla, P., Kapoor, A., Lan, L., Mehta, S., Mithani, R., Panozzo, C. A., Simorellis, A. K., Kuter, B. J., Schödel, F., Huang, W., Reuter, C., Slobod, K., Stoszek, S. K., Shaw, C. A., Miller, J. M., Das, R. & Chen, G. L. Efficacy and Safety of an mRNA-Based RSV PreF Vaccine in Older Adults. New England Journal of Medicine 389, 2233–2244. DOI: 10.1056/NEJMoa2307079 (Dec. 13, 2023).

6. Moderna Receives U.S. FDA Approval for RSV Vaccine mRESVIA(R) https://investors.modernatx.com/news/news-details/2024/Moderna-Receives-U.S.-FDA-Approval-for-RSV-Vaccine-mRESVIAR/default.aspx (2025).

7. Sayour, E. J., Boczkowski, D., Mitchell, D. A. & Nair, S. K. Cancer mRNA vaccines: clinical advances and future opportunities. Nature Reviews Clinical Oncology 21, 489–500. DOI: 10.1038/s41571-024-00902-1 (July 2024).

8. Verbeke, R., Hogan, M. J., Loré, K. & Pardi, N. Innate immune mechanisms of mRNA vaccines. Immunity 55, 1993–2005. DOI: 10.1016/j.immuni.2022.10.014 (Nov. 2022).

9. Jeong, M., Lee, Y., Park, J., Jung, H. & Lee, H. Lipid nanoparticles (LNPs) for in vivo RNA delivery and their breakthrough technology for future applications. Advanced Drug Delivery Reviews 200, 114990. DOI: 10.1016/j.addr.2023.114990 (Sept. 1, 2023).

10. Zwolsman, R., Darwish, Y. B., Kluza, E. & van der Meel, R. Engineering Lipid Nanoparticles for mRNA Immunotherapy. Wiley Interdisciplinary Reviews. Nanomedicine and Nanobiotechnology 17, e70007. DOI: 10.1002/wnan.70007 (2025).

11. Hamilton, A. G., Swingle, K. L., Thatte, A. S., Mukalel, A. J., Safford, H. C., Billingsley, M. M., El-Mayta, R. D., Han, X., Nachod, B. E., Joseph, R. A., Metzloff, A. E. & Mitchell, M. J. High-Throughput In Vivo Screening Identifies Differential Influences on mRNA Lipid Nanoparticle Immune Cell Delivery by Administration Route. ACS Nano 18, 16151–16165. DOI: 10.1021/acsnano.4c01171 (June 2024).

12. Dahlman, J. E., Kauffman, K. J., Xing, Y., Shaw, T. E., Mir, F. F., Dlott, C. C., Langer, R., Anderson, D. G. & Wang, E. T. Barcoded nanoparticles for high throughput in vivo discovery of targeted therapeutics. Proceedings of the National Academy of Sciences 114, 2060–2065. DOI: 10.1073/pnas.1620874114 (Feb. 2017).

13. El-Mayta, R., Zhang, R., Shepherd, S. J., Wang, F., Billingsley, M. M., Dudkin, V., Klein, D., Lu, H. D. & Mitchell, M. J. A Nanoparticle Platform for Accelerated In Vivo Oral Delivery Screening of Nucleic Acids. Advanced Therapeutics 4, 2000111. DOI: 10.1002/adtp.202000111 (2021).

14. Lokugamage, M. P., Sago, C. D., Gan, Z., Krupzak, B. & Dahlman, J. E. Constrained nanoparticles deliver siRNA and sgRNA to T cells in vivo without targeting ligands. Advanced Materials 31, e1902251. DOI: 10.1002/adma.201902251 (Oct. 2019).

15. Eygeris, Y., Gupta, M., Kim, J., Jozic, A., Gautam, M., Renner, J., Nelson, D., Bloom, E., Tuttle, A., Stoddard, J., Reynaga, R., Neuringer, M., Lauer, A. K., Ryals, R. C. & Sahay, G. Thiophene-based lipids for mRNA delivery to pulmonary and retinal tissues. Proceedings of the National Academy of Sciences 121, e2307813120. DOI: 10.1073/pnas.2307813120 (Mar. 12, 2024).

16. Guimaraes, P. P. G., Zhang, R., Spektor, R., Tan, M., Chung, A., Billingsley, M. M., El-Mayta, R., Riley, R. S., Wang, L., Wilson, J. M. & Mitchell, M. J. Ionizable lipid nanoparticles encapsulating barcoded mRNA for accelerated in vivo delivery screening. Journal of Controlled Release 316, 404–417. DOI: 10.1016/j.jconrel.2019.10.028 (Dec. 2019).

17. Zhang, R., El-Mayta, R., Murdoch, T. J., Warzecha, C. C., Billingsley, M. M., Shepherd, S. J., Gong, N., Wang, L., Wilson, J. M., Lee, D. & Mitchell, M. J. Helper lipid structure influences protein adsorption and delivery of lipid nanoparticles to spleen and liver. Biomaterials Science 9, 1449–1463. DOI: 10.1039/D0BM01609H (2021).

18. Akinc, A., Querbes, W., De, S., Qin, J., Frank-Kamenetsky, M., Jayaprakash, K. N., Jayaraman, M., Rajeev, K. G., Cantley, W. L., Dorkin, J. R., Butler, J. S., Qin, L., Racie, T., Sprague, A., Fava, E., Zeigerer, A., Hope, M. J., Zerial, M., Sah, D. W., Fitzgerald, K., Tracy, M. A., Manoharan, M., Koteliansky, V., Fougerolles, A. d. & Maier, M. A. Targeted Delivery of RNAi Therapeutics With Endogenous and Exogenous Ligand-Based Mechanisms. Molecular Therapy 18, 1357–1364. DOI: 10.1038/mt.2010.85 (July 2010).

19. Suzuki, Y. & Ishihara, H. Structure, activity and uptake mechanism of siRNA-lipid nanoparticles with an asymmetric ionizable lipid. International Journal of Pharmaceutics 510, 350–358. DOI: 10.1016/j.ijpharm.2016.06.124 (Aug. 2016).

20. Francia, V., Schiffelers, R. M., Cullis, P. R. & Witzigmann, D. The Biomolecular Corona of Lipid Nanoparticles for Gene Therapy. Bioconjugate Chemistry 31, 2046–2059. DOI: 10.1021/acs.bioconjchem.0c00366 (Sept. 2020).

21. Sato, Y., Kinami, Y., Hashiba, K. & Harashima, H. Different kinetics for the hepatic uptake of lipid nanoparticles between the apolipoprotein E/low density lipoprotein receptor and the N -acetyl-d-galactosamine/asialoglycoprotein receptor pathway. Journal of Controlled Release 322, 217–226. DOI: 10.1016/j.jconrel.2020.03.006 (June 2020).

22. Dilliard, S. A., Cheng, Q. & Siegwart, D. J. On the mechanism of tissue-specific mRNA delivery by selective organ targeting nanoparticles. Proceedings of the National Academy of Sciences 118, e2109256118. DOI: 10.1073/pnas.2109256118 (Dec. 2021).

23. Kauffman, K. J., Dorkin, J. R., Yang, J. H., Heartlein, M. W., DeRosa, F., Mir, F. F., Fenton, O. S. & Anderson, D. G. Optimization of Lipid Nanoparticle Formulations for mRNA Delivery in Vivo with Fractional Factorial and Definitive Screening Designs. Nano Letters 15, 7300–7306. DOI: 10.1021/acs.nanolett.5b02497 (Nov. 2015).

24. Ball, R. L., Hajj, K. A., Vizelman, J., Bajaj, P. & Whitehead, K. A. Lipid Nanoparticle Formulations for Enhanced Co-delivery of siRNA and mRNA. Nano Letters 18, 3814–3822. DOI: 10.1021/acs.nanolett.8b01101 (June 2018).

25. Hamilton, A. G., Swingle, K. L., Joseph, R. A., Mai, D., Gong, N., Billingsley, M. M., Alameh, M.-G., Weissman, D., Sheppard, N. C., June, C. H. & Mitchell, M. J. Ionizable Lipid Nanoparticles with Integrated Immune Checkpoint Inhibition for mRNA CAR T Cell Engineering. Advanced Healthcare Materials 12, 2301515. DOI: 10.1002/adhm.202301515 (Aug. 2023).

26. Sago, C. D., Lokugamage, M. P., Paunovska, K., Vanover, D. A., Monaco, C. M., Shah, N. N., Gamboa Castro, M., Anderson, S. E., Rudoltz, T. G., Lando, G. N., Munnilal Tiwari, P., Kirschman, J. L., Willett, N., Jang, Y. C., Santangelo, P. J., Bryksin, A. V. & Dahlman, J. E. High-throughput in vivo screen of functional mRNA delivery identifies nanoparticles for endothelial cell gene editing. Proceedings of the National Academy of Sciences 115, E9944–E9952. DOI: 10.1073/pnas.1811276115 (Oct. 2018).

27. Buschmann, T. & Bystrykh, L. V. Levenshtein error-correcting barcodes for multiplexed DNA sequencing. BMC Bioinformatics 14, 272. DOI: 10.1186/1471-2105-14-272 (Sept. 2013).

28. Houseley, J. & Tollervey, D. The Many Pathways of RNA Degradation. Cell 136, 763–776. DOI:10.1016/j.cell.2009.01.019 (Feb. 2009).

29. Glisovic, T., Bachorik, J. L., Yong, J. & Dreyfuss, G. RNA-binding proteins and post-transcriptional gene regulation. FEBS Letters. Nuclear Dynamics and Cytoskeleton Signaling 582, 1977–1986. DOI: 10.1016/j.febslet.2008.03.004 (June 2008).

30. Denzler, R., McGeary, S. E., Title, A. C., Agarwal, V., Bartel, D. P. & Stoffel, M. Impact of MicroRNA Levels, Target-Site Complementarity, and Cooperativity on Competing Endogenous RNA-Regulated Gene Expression. Molecular Cell 64, 565–579. DOI: 10.1016/j.molcel.2016.09.027 (Nov. 2016).

31. Landgraf, P., Rusu, M., Sheridan, R., Sewer, A., Iovino, N., Aravin, A., Pfeffer, S., Rice, A., Kamphorst, A. O., Landthaler, M., Lin, C., Socci, N. D., Hermida, L., Fulci, V., Chiaretti, S., Foà, R., Schliwka, J., Fuchs, U., Novosel, A., Müller, R.-U., Schermer, B., Bissels, U., Inman, J., Phan, Q., Chien, M., Weir, D. B., Choksi, R., De Vita, G., Frezzetti, D., Trompeter, H.-I., Hornung, V., Teng, G., Hartmann, G., Palkovits, M., Di Lauro, R., Wernet, P., Macino, G., Rogler, C. E., Nagle, J. W., Ju, J., Papavasiliou, F. N., Benzing, T., Lichter, P., Tam, W., Brownstein, M. J., Bosio, A., Borkhardt, A., Russo, J. J., Sander, C., Zavolan, M. & Tuschl, T. A Mammalian microRNA Expression Atlas Based on Small RNA Library Sequencing. Cell 129, 1401–1414. DOI: 10.1016/j.cell.2007.04.040 (June 2007).

32. Ludwig, N., Leidinger, P., Becker, K., Backes, C., Fehlmann, T., Pallasch, C., Rheinheimer, S., Meder, B., Stähler, C., Meese, E. & Keller, A. Distribution of miRNA expression across human tissues. Nucleic Acids Research 44, 3865–3877. DOI: 10.1093/nar/gkw116 (May 2016).

33. An integrated expression atlas of miRNAs and their promoters in human and mouse. Nature Biotechnology 35. Publisher: Nature Publishing Group, 872–878. DOI: 10.1038/nbt.3947 (Sept. 2017).

34. Han, X., Xu, Y., Ricciardi, A., Xu, J., Palanki, R., Chowdhary, V., Xue, L., Gong, N., Alameh, M.-G., Peranteau, W. H., Wilson, J. M., Weissman, D. & Mitchell, M. J. Plug-and-play assembly of biodegradable ionizable lipids for potent mRNA delivery and gene editing in vivo. Preprint at https://www.biorxiv.org/content/10.1101/2025.02.25.640222v1 (mMar. 2025).

35. Love, K. T., Mahon, K. P., Levins, C. G., Whitehead, K. A., Querbes, W., Dorkin, J. R., Qin, J., Cantley, W., Qin, L. L., Racie, T., Frank-Kamenetsky, M., Yip, K. N., Alvarez, R., Sah, D. W. Y., de Fougerolles, A., Fitzgerald, K., Koteliansky, V., Akinc, A., Langer, R. & Anderson, D. G. Lipid-like materials for low-dose, in vivo gene silencing. Proceedings of the National Academy of Sciences 107, 1864–1869. DOI: 10.1073/pnas.0910603106 (Feb. 2010).

36. Dong, Y., Love, K. T., Dorkin, J. R., Sirirungruang, S., Zhang, Y., Chen, D., Bogorad, R. L., Yin, H., Chen, Y., Vegas, A. J., Alabi, C. A., Sahay, G., Olejnik, K. T., Wang, W., Schroeder, A., Lytton-Jean, A. K. R., Siegwart, D. J., Akinc, A., Barnes, C., Barros, S. A., Carioto, M., Fitzgerald, K., Hettinger, J., Kumar, V., Novobrantseva, T. I., Qin, J., Querbes, W., Koteliansky, V., Langer, R. & Anderson, D. G. Lipopeptide nanoparticles for potent and selective siRNA delivery in rodents and nonhuman primates. Proceedings of the National Academy of Sciences 111, 3955–3960 (Mar. 2014).

37. Jayaraman, M., Ansell, S. M., Mui, B. L., Tam, Y. K., Chen, J., Du, X., Butler, D., Eltepu, L., Matsuda, S., Narayanannair, J. K., Rajeev, K. G., Hafez, I. M., Akinc, A., Maier, M. A., Tracy, M. A., Cullis, P. R., Madden, T. D., Manoharan, M. & Hope, M. J. Maximizing the potency of siRNA lipid nanoparticles for hepatic gene silencing in vivo. Angewandte Chemie (International Ed. in English) 51, 8529–8533. DOI: 10.1002/anie.201203263 (Aug. 2012).

38. Hajj, K. A., Ball, R. L., Deluty, S. B., Singh, S. R., Strelkova, D., Knapp, C. M. & Whitehead, K. A. Branched-Tail Lipid Nanoparticles Potently Deliver mRNA In Vivo due to Enhanced Ionization at Endosomal pH. Small 15, 1805097. DOI: 10.1002/smll.201805097 (2019).

39. Sabnis, S., Kumarasinghe, E. S., Salerno, T., Mihai, C., Ketova, T., Senn, J. J., Lynn, A., Bulychev, A., McFadyen, I., Chan, J., Almarsson, Ö., Stanton, M. G. & Benenato, K. E. A Novel Amino Lipid Series for mRNA Delivery: Improved Endosomal Escape and Sustained Pharmacology and Safety in Non-human Primates. Molecular Therapy 26, 1509–1519. DOI: 10.1016/j.ymthe.2018.03.010 (June 2018).

40. Verbeke, R., Lentacker, I., De Smedt, S. C. & Dewitte, H. The dawn of mRNA vaccines: The COVID-19 case. Journal of Controlled Release 333, 511–520. DOI: 10.1016/j.jconrel.2021.03.043 (May 2021).

41. Thatte, A. S., Hamilton, A. G., Nachod, B. E., Mukalel, A. J., Billingsley, M. M., Palanki, R., Swingle, K. L. & Mitchell, M. J. mRNA Lipid Nanoparticles for Ex Vivo Engineering of Immunosuppressive T Cells for Autoimmunity Therapies. Nano Letters 23, 10179–10188. DOI: 10.1021/acs.nanolett.3c02573 (Nov. 2023).

42. Mukalel, A. J., Hamilton, A. G., Billingsley, M. M., Li, J., Thatte, A. S., Han, X., Safford, H. C., Padilla, M. S., Papp, T., Parhiz, H., Weissman, D. & Mitchell, M. J. Oxidized mRNA Lipid Nanoparticles for In Situ Chimeric Antigen Receptor Monocyte Engineering. Advanced Functional Materials 34, 2312038. DOI: 10.1002/adfm.202312038 (July 2024).

43. Swingle, K. L., Hamilton, A. G., Safford, H. C., Geisler, H. C., Thatte, A. S., Palanki, R., Murray, A. M., Han, E. L., Mukalel, A. J., Han, X., Joseph, R. A., Ghalsasi, A. A., Alameh, M.-G., Weissman, D. & Mitchell, M. J. Placenta-tropic VEGF mRNA lipid nanoparticles ameliorate murine pre-eclampsia. Nature, 1–10. DOI: 10.1038/s41586-024-08291-2 (Dec. 2024).

44. Xu, X., Wang, X., Liao, Y.-P., Luo, L., Xia, T. & Nel, A. E. Use of a Liver-Targeting Immune-Tolerogenic mRNA Lipid Nanoparticle Platform to Treat Peanut-Induced Anaphylaxis by Single- and Multiple-Epitope Nucleotide Sequence Delivery. ACS Nano 17, 4942–4957. DOI: 10.1021/acsnano.2c12420 (Mar. 2023).

45. Bashiri, G., Padilla, M. S., Swingle, K. L., Shepherd, S. J., Mitchell, M. J. & Wang, K. Nanoparticle protein corona: from structure and function to therapeutic targeting. Lab on a Chip 23, 1432–1466. DOI: 10.1039/d2lc00799a (Jan. 2023).

46. Su, A. I., Cooke, M. P., Ching, K. A., Hakak, Y., Walker, J. R., Wiltshire, T., Orth, A. P., Vega, R. G., Sapinoso, L. M., Moqrich, A., Patapoutian, A., Hampton, G. M., Schultz, P. G. & Hogenesch, J. B. Large-scale analysis of the human and mouse transcriptomes. Proceedings of the National Academy of Sciences of the United States of America 99, 4465–4470. DOI: 10.1073/pnas.012025199 (Apr. 2002).

47. Su, A. I., Wiltshire, T., Batalov, S., Lapp, H., Ching, K. A., Block, D., Zhang, J., Soden, R., Hayakawa, M., Kreiman, G., Cooke, M. P., Walker, J. R. & Hogenesch, J. B. A gene atlas of the mouse and human protein-encoding transcriptomes. Proceedings of the National Academy of Sciences of the United States of America 101, 6062–6067. DOI: 10.1073/pnas.0400782101 (Apr. 2004).

48. Paunovska, K., Da Silva Sanchez, A. J., Lokugamage, M. P., Loughrey, D., Echeverri, E. S., Cristian, A., Hatit, M. Z. C., Santangelo, P. J., Zhao, K. & Dahlman, J. E. The Extent to Which Lipid Nanoparticles Require Apolipoprotein E and Low-Density Lipoprotein Receptor for Delivery Changes with Ionizable Lipid Structure. Nano Letters 22, 10025–10033. DOI: 10.1021/acs.nanolett.2c03741 (Dec. 2022).

49. Chen, D., Love, K. T., Chen, Y., Eltoukhy, A. A., Kastrup, C., Sahay, G., Jeon, A., Dong, Y., Whitehead, K. A. & Anderson, D. G. Rapid Discovery of Potent siRNA-Containing Lipid Nanoparticles Enabled by Controlled Microfluidic Formulation. Journal of the American Chemical Society 134, 6948–6951. DOI: 10.1021/ja301621z (Apr. 2012).

50. R Core Team. R: A Language and Environment for Statistical Computing R Foundation for Statistical Computing (Vienna, Austria, 2023).

51. Rudolph, K. box: Write Reusable, Composable and Modular R Code (2024).

52. Wickham, H., François, R., Henry, L., Müller, K. & Vaughan, D. dplyr: A Grammar of Data Manipulation (2023).

53. Wickham, H. forcats: Tools for Working with Categorical Variables (Factors) (2023).

54. Clarke, E. & Sherrill-Mix, S. ggbeeswarm: Categorical Scatter (Violin Point) Plots (2017).

55. Wickham, H. ggplot2: Elegant Graphics for Data Analysis (Springer-Verlag New York, 2016).

56. Wilke, C. O. & Wiernik, B. M. ggtext: Improved Text Rendering Support for ‘ggplot2’ (2022).

57. Pedersen, T. L. patchwork: The Composer of Plots (2024).

58. Ooms, J. pdftools: Text Extraction, Rendering and Converting of PDF Documents (2023).

59. Wickham, H., Hester, J. & Bryan, J. readr: Read Rectangular Text Data (2024).

60. Ooms, J. rsvg: Render SVG Images into PDF, PNG, (Encapsulated) PostScript, or Bitmap Arrays (2023).

61. Wickham, H., Henry, L., Pedersen, T. L., Luciani, T. J., Decorde, M. & Lise, V. svglite: An ‘SVG’ Graphics Device (2023).

62. Wickham, H., Vaughan, D. & Girlich, M. tidyr: Tidy Messy Data (2024).

63. Smith, T., Heger, A. & Sudbery, I. UMI-tools: modeling sequencing errors in Unique Molecular Identifiers to improve quantification accuracy. Genome Research 27, 491–499. DOI: 10.1101/gr.209601.116 (Mar. 2017).

64. Benjamini, Y. & Hochberg, Y. Controlling the False Discovery Rate: A Practical and Powerful Approach to Multiple Testing. Journal of the Royal Statistical Society. Series B (Methodological) 57, 289–300 (1995).

65. Piao, X., Yadav, V., Wang, E., Chang, W., Tau, L., Lindenmuth, B. E. & Wang, S. X. Double-stranded RNA reduction by chaotropic agents during in vitro transcription of messenger RNA. Molecular Therapy - Nucleic Acids 29, 618–624. DOI: 10.1016/j.omtn.2022.08.001 (Sept. 2022).

66. Welsh, J. A., Pol, E. V. D., Arkesteijn, G. J., Bremer, M., Brisson, A., Coumans, F., Dignat-George, F., Duggan, E., Ghiran, I., Giebel, B., Görgens, A., Hendrix, A., Lacroix, R., Lannigan, J., Libregts, S. F., Lozano-Andrés, E., Morales-Kastresana, A., Robert, S., Rond, L. D., Tertel, T., Tigges, J., Wever, O. D., Yan, X., Nieuwland, R., Wauben, M. H., Nolan, J. P. & Jones, J. C. MIFlowCyt-EV: a framework for standardized reporting of extracellular vesicle flow cytometry experiments. Journal of Extracellular Vesicles 9, 1713526. DOI: 10.1080/20013078.2020.1713526 (Feb. 2020).

67. Han, X., Alameh, M.-G., Butowska, K., Knox, J. J., Lundgreen, K., Ghattas, M., Gong, N., Xue, L., Xu, Y., Lavertu, M., Bates, P., Xu, J., Nie, G., Zhong, Y., Weissman, D. & Mitchell, M. J. Adjuvant lipidoid-substituted lipid nanoparticles augment the immunogenicity of SARS-CoV-2 mRNA vaccines. Nature Nanotechnology 18, 1105–1114. DOI: 10.1038/s41565-023-01404-4 (Sept. 2023).

68. Frey, A., Di Canzio, J. & Zurakowski, D. A statistically defined endpoint titer determination method for immunoassays. Journal of Immunological Methods 221, 35–41. DOI: 10.1016/S0022-1759(98)00170-7 (Dec. 1, 1998).

69. Robinson, D., Hayes, A. & Couch, S. broom: Convert Statistical Objects into Tidy Tibbles (2024).

70. Scott, J. ggborderline: Line Plots that Pop (2022).

71. Grolemund, G. & Wickham, H. Dates and Times Made Easy with lubridate. Journal of Statistical Software 40, 1–25 (2011).

72. Ooms, J. magick: Advanced Graphics and Image-Processing in R (2024).

73. Kassambara, A., Kosinski, M. & Biecek, P. survminer: Drawing Survival Curves using ‘ggplot2’ (2021).

74. Therneau, T. M. A Package for Survival Analysis in R (2024).

75. Dolstra, E. The purely functional software deployment model (Jan. 2006).

76. Terry M. Therneau & Patricia M. Grambsch. Modeling Survival Data: Extending the Cox Model (Springer, New York, 2000).

