## Supplementary information for "High-throughput *in vivo* screening using barcoded mRNA identifies lipid nanoparticles with extrahepatic tropism for cancer immunotherapy"

| Barcode | Sequence |
|---|---|
| 1 | CCTCTTGTGTGG |
| 2 | AATGTTCTCTCC |
| 3 | GGAGAAGAAGAA |
| 4 | TATGGTTGTGTG |
| 5 | GGTCGGTGTTGT |
| 6 | CCTTCTGTTCTG |
| 7 | CTCTTCGCCTCT |
| 8 | TTCGGTTCTTCC |
| 9 | TTATCCTCTCCA |
| 10 | GTGTTCTCCTTA |
| 11 | TTCTCGGTGTGG |
| 12 | TTGTTGGCTGCC |
| 13 | CTTGGTGGTTCC |
| 14 | GGTCCTTCCTCA |
| 15 | TGGTGTGGTCTC |
| 16 | GTGGTTGGTTAA |
| 17 | GTGTTGTGTCCG |
| 18 | AACTCCTTCCGG |
| 19 | AATCCATCTTCC |
| 20 | TTCACACCACAA |
| 21 | AAGGAAGGTGGA |
| 22 | AAGTGGTTGCGG |
| 23 | CCAACACCAGCA |
| 24 | CCAAGGTGTGTG |
| 25 | GAGATTGTGTTG |
| 26 | GCGCCTTCTCTA |
| 27 | TCCTCCGCTTGT |
| 28 | CTTCCTTGTCGG |
| 29 | AATTATTGCCGG |
| 30 | CCAATTCTCCTA |
| 31 | TTATGATGGTGG |
| 32 | TGTGTCTTCGCC |
| 33 | GGAGTCTCTCTC |
| 34 | ACCTTCCTCTAA |
| 35 | AACCTCTCTCAA |
| 36 | TAGAGAGGAGAA |
| 37 | CCACACATTCAA |
| 38 | TAATCCTCCGCG |
| 39 | AGGCCACACCAA |
| 40 | GGCAATCCTCTT |
| 41 | ACAACCAATCCA |
| 42 | GACCACACATAA |
| 43 | TTGTGCGCGTTC |
| 44 | TCCGTTGTCTCC |
| 45 | CACCACTCATCC |
| 46 | AGAATAGGTTGG |
| 47 | TTCCTTCGTTGA |
| 48 | TGTTGTTGGAGA |
| 49 | CGGTTGTCCTGT |

| Barcode | Sequence |
|---|---|
| 50 | GGAGGAACACAA |
| 51 | TCCTCTTCACCG |
| 52 | ATAGTGGCGTGG |
| 53 | AAGAGGAGCGAA |
| 54 | AGGAAGAGGCAA |
| 55 | ATTGTGTGATGG |
| 56 | AACCACCGGCAA |
| 57 | ACACACAACCGA |
| 58 | TTCCGCCACTCA |
| 59 | CTTGTGGTGTAA |
| 60 | ACGCTGTTGTGG |
| 61 | AAGAACAAGCGG |
| 62 | TGGCTCTTCTCA |
| 63 | CTCCAACCTTAA |
| 64 | TGGTGGCGCTTA |
| 65 | ACAAGAGAGGTA |
| 66 | TTGGTGTATTGC |
| 67 | GGCCACCTTATA |
| 68 | AGGTGTTCTTGA |
| 69 | ACCTCTCTTGGA |
| 70 | TTATAGGCTTGG |
| 71 | TCCGTCTTGTTA |
| 72 | GCCATCTCCTCA |
| 73 | GGTTGTTGCTAA |
| 74 | CCTCCGTTCCAA |
| 75 | AACCAAGCCTAA |
| 76 | TCGTTCTGTGTG |
| 77 | CCGCGTTATTAA |
| 78 | AAGCAACCTCCA |
| 79 | CACTTCATTCTC |
| 80 | GTACAACACCAA |
| 81 | TGCCTCCATCCA |
| 82 | TAGAGGATGTGG |
| 83 | TTCTCCAATACC |
| 84 | GATTGCTTCTTG |
| 85 | AACACCGCTTAA |
| 86 | TGTGGAAGAGAA |
| 87 | ACACCGCTCTGT |
| 88 | GCCGCTTATTCC |
| 89 | GTCTCTTGGTGC |
| 90 | CTCGCTTATCTT |
| 91 | CATACTTCCTTG |
| 92 | TTCTTCCTGCAA |
| 93 | ATTACCTCGCGG |
| 94 | GGAGAGAATTGG |
| 95 | CATCGTGGTTGG |
| 96 | GGTGAGAGGTAA |
| 97 | AATAGGTAGTGG |
| 98 | CACCTAACACAA |

| Barcode | Sequence |
|---|---|
| 99 | TGGAACAACCAA |
| 100 | ACCACGGCTGTT |
| 101 | TCCAATCTTCTG |
| 102 | TCCTTCTACCTG |
| 103 | CCTTCTAACACC |
| 104 | GGTCTTGATCTT |
| 105 | AAGGCTCCTCAC |
| 106 | TGGAGGTAGTTG |
| 107 | GTGGAGTCTTCC |
| 108 | AGGTCACCTCTT |
| 109 | ACGGTGTGTTAA |
| 110 | CCTCACCACTTG |
| 111 | GTAACACTCTTC |
| 112 | GCCAATGGTTGG |
| 113 | GTTGGCCTTGTA |
| 114 | TAACACAGCCAA |
| 115 | CGTTCCTCTTGA |
| 116 | GTCCTCTGTTCG |
| 117 | TCTCCACTCTAA |
| 118 | TACACTTGTTGG |
| 119 | CTTCTTACGCGG |
| 120 | ACCAGCCACCTA |
| 121 | GGTTGCTTATCC |
| 122 | AAGAACCACTTG |
| 123 | AGAGAGAGCCTA |
| 124 | TTGTGGTAGCGC |
| 125 | CCTCATTATTCG |
| 126 | TTACGCCTTCGC |
| 127 | ATACCTGTGTGG |
| 128 | ACCAACCTTGCA |
| 129 | CAAGGCCTTCCA |
| 130 | TCCATCCTATTC |
| 131 | ATTGGCGTCTGG |
| 132 | TCCTATTCGTGG |
| 133 | GAGTGGTGGTCA |
| 134 | CGGAAGAAGGTG |
| 135 | GGTTGGAGTGAA |
| 136 | CCGACTCTCTCA |
| 137 | CGCCACAACTCA |
| 138 | TGTGGCCTCCAA |
| 139 | GCCGAACAACAA |
| 140 | ACAGACTCTTCC |
| 141 | AACTGCTGGTGG |
| 142 | CATATATGGCGG |
| 143 | TCTGCTTCTCGG |
| 144 | GCTTACCTTCTA |
| 145 | CCGTTATATACC |
| 146 | CCTTGTTGTCAA |
| 147 | ACAACGCGCCAA |

| Barcode | Sequence |
|---|---|
| 148 | CAAGAACACTCA |
| 149 | GCTGTTCGTTCG |
| 150 | TCCACCTCCGAA |
| 151 | GTTGGAGGCGTA |
| 152 | GGTTATCTGCGG |
| 153 | TAGGAGCCTCCT |
| 154 | TCCAGGAGAGAA |
| 155 | ACACAGAGACAA |
| 156 | CCGGAGAATTAA |
| 157 | AACAGGAGGCTC |
| 158 | TGTTCAGTTCTC |
| 159 | GGTGGTCTATAA |
| 160 | GGTGTTCAGTGG |
| 161 | GGCATTGTTCTG |
| 162 | CAACCGTCGCTT |
| 163 | CAATAACCTTCG |
| 164 | TTCTTGGAGGAA |
| 165 | CGGTGGATGGAA |
| 166 | TATAGGAGACGG |
| 167 | CTTCCTACTCAA |
| 168 | ATGCCTCTTCGG |
| 169 | GAAGTGAGAGAA |
| 170 | AGAACGACACAA |
| 171 | AACGGCACACAA |
| 172 | CTTCTCCAGGTT |
| 173 | AGGACGGTTCTC |
| 174 | TATTCTGCCTCA |
| 175 | TCTGGTCTGCTT |
| 176 | GGAAGTTGTGAA |
| 177 | ACCTTGTATTCC |
| 178 | GAACACAAGGAA |
| 179 | TGGAACTGTGTT |
| 180 | CAAGAGAAGGCC |
| 181 | CTTGTTCATTGC |
| 182 | CACACCTCTGCA |
| 183 | TTCTCGTGCTTA |
| 184 | ACTTGCCTTCGG |
| 185 | CTTCCACTGTTA |
| 186 | AACAAGCGAGAA |
| 87 | ACGTGAGGTTGG |
| 188 | GGCACAACAGAA |
| 189 | AGAACACCTCGA |
| 190 | AATTAAGCTCCG |
| 191 | TAACACCGGTCG |
| 192 | AGGATGAGTGTG |
| 193 | GGAGTGGATATA |
| 194 | TAATGGAGGTGA |
| 195 | GGAAGGCATTAT |
| 196 | AGGTTATGGTGA |

| Barcode | Sequence |
| --- | --- |
| 197 | AGTATTCTCTGG |
| 198 | TATCCATTGTGG |
| 199 | CTTGTTCCACAA |
| 200 | CGAGCGGTGTTA |

**Supplementary Table 1:** Top 200 barcode sequences used to produce an initial library of b-mRNAs for *in vivo* evaluation.

| Barcode | Sequence | Index |
|---|---|---|
| 4 | TATGGTTGTGTG | 1 |
| 5 | GGTCGGTGTTGT | 2 |
| 6 | CCTTCTGTTCTG | 3 |
| 7 | CTCTTCGCCTCT | 4 |
| 8 | TTCGGTTCTTCC | 5 |
| 9 | TTATCCTCTCCA | 6 |
| 10 | GTGTTCTCCTTA | 7 |
| 13 | CTTGGTGGTTCC | 8 |
| 14 | GGTCCTTCCTCA | 9 |
| 15 | TGGTGTGGTCTC | 10 |
| 17 | GTGTTGTGTCCG | 11 |
| 18 | AACTCCTTCCGG | 12 |
| 20 | TTCACACCACAA | 13 |
| 21 | AAGGAAGGTGGA | 14 |
| 23 | CCAACACCAGCA | 15 |
| 24 | CCAAGGTGTGTG | 16 |
| 25 | GAGATTGTGTTG | 17 |
| 26 | GCGCCTTCTCTA | 18 |
| 27 | TCCTCCGCTTGT | 19 |
| 29 | AATTATTGCCGG | 20 |
| 32 | TGTGTCTTCGCC | 21 |
| 39 | AGGCCACACCAA | 22 |
| 41 | ACAACCAATCCA | 23 |
| 44 | TCCGTTGTCTCC | 24 |
| 46 | AGAATAGGTTGG | 25 |
| 47 | TTCCTTCGTTGA | 26 |
| 49 | CGGTTGTCCTGT | 27 |
| 50 | GGAGGAACACAA | 28 |
| 51 | TCCTCTTCACCG | 29 |
| 53 | AAGAGGAGCGAA | 30 |
| 55 | ATTGTGTGATGG | 31 |
| 56 | AACCACCGGCAA | 32 |
| 57 | ACACACAACCGA | 33 |
| 58 | TTCCGCCACTCA | 34 |
| 59 | CTTGTGGTGTAA | 35 |
| 61 | AAGAACAAGCGG | 36 |
| 62 | TGGCTCTTCTCA | 37 |
| 64 | TGGTGGCGCTTA | 38 |
| 65 | ACAAGAGAGGTA | 39 |
| 66 | TTGGTGTATTGC | 40 |
| 68 | AGGTGTTCTTGA | 41 |
| 69 | ACCTCTCTTGGA | 42 |
| 71 | TCCGTCTTGTTA | 43 |
| 72 | GCCATCTCCTCA | 44 |
| 73 | GGTTGTTGCTAA | 45 |
| 74 | CCTCCGTTCCAA | 46 |
| 75 | AACCAAGCCTAA | 47 |
| 78 | AAGCAACCTCCA | 48 |
| 79 | CACTTCATTCTC | 49 |

| Barcode | Sequence | Index |
|---|---|---|
| 80 | GTACAACACCAA | 50 |
| 81 | TGCCTCCATCCA | 51 |
| 84 | GATTGCTTCTTG | 52 |
| 87 | ACACCGCTCTGT | 53 |
| 89 | GTCTCTTGGTGC | 54 |
| 90 | CTCGCTTATCTT | 55 |
| 91 | CATACTTCCTTG | 56 |
| 94 | GGAGAGAATTGG | 57 |
| 96 | GGTGAGAGGTAA | 58 |
| 97 | AATAGGTAGTGG | 59 |
| 98 | CACCTAACACAA | 60 |
| 99 | TGGAACAACCAA | 61 |
| 100 | ACCACGGCTGTT | 62 |
| 101 | TCCAATCTTCTG | 63 |
| 102 | TCCTTCTACCTG | 64 |
| 103 | CCTTCTAACACC | 65 |
| 104 | GGTCTTGATCTT | 66 |
| 105 | AAGGCTCCTCAC | 67 |
| 106 | TGGAGGTAGTTG | 68 |
| 107 | GTGGAGTCTTCC | 69 |
| 108 | AGGTCACCTCTT | 70 |
| 109 | ACGGTGTGTTAA | 71 |
| 110 | CCTCACCACTTG | 72 |
| 111 | GTAACACTCTTC | 73 |
| 112 | GCCAATGGTTGG | 74 |
| 113 | GTTGGCCTTGTA | 75 |
| 114 | TAACACAGCCAA | 76 |
| 115 | CGTTCCTCTTGA | 77 |
| 116 | GTCCTCTGTTCG | 78 |
| 118 | TACACTTGTTGG | 79 |
| 119 | CTTCTTACGCGG | 80 |
| 120 | ACCAGCCACCTA | 81 |
| 121 | GGTTGCTTATCC | 82 |
| 122 | AAGAACCACTTG | 83 |
| 123 | AGAGAGAGCCTA | 84 |
| 124 | TTGTGGTAGCGC | 85 |
| 125 | CCTCATTATTCG | 86 |
| 127 | ATACCTGTGTGG | 87 |
| 128 | ACCAACCTTGCA | 88 |
| 129 | CAAGGCCTTCCA | 89 |
| 130 | TCCATCCTATTC | 90 |
| 131 | ATTGGCGTCTGG | 91 |
| 133 | GAGTGGTGGTCA | 92 |
| 135 | GGTTGGAGTGAA | 93 |
| 136 | CCGACTCTCTCA | 94 |
| 138 | TGTGGCCTCCAA | 95 |
| 140 | ACAGACTCTTCC | 96 |
| 142 | CATATATGGCGG | 97 |
| 143 | TCTGCTTCTCGG | 98 |

| Barcode | Sequence | Index |
|---|---|---|
| 146 | CCTTGTTGTCAA | 99 |
| 147 | ACAACGCGCCAA | 100 |
| 149 | GCTGTTCGTTCG | 101 |
| 150 | TCCACCTCCGAA | 102 |
| 151 | GTTGGAGGCGTA | 103 |
| 153 | TAGGAGCCTCCT | 104 |
| 155 | ACACAGAGACAA | 105 |
| 157 | AACAGGAGGCTC | 106 |
| 158 | TGTTCAGTTCTC | 107 |
| 161 | GGCATTGTTCTG | 108 |
| 162 | CAACCGTCGCTT | 109 |
| 163 | CAATAACCTTCG | 110 |
| 165 | CGGTGGATGGAA | 111 |
| 166 | TATAGGAGACGG | 112 |
| 167 | CTTCCTACTCAA | 113 |
| 169 | GAAGTGAGAGAA | 114 |
| 171 | AACGGCACACAA | 115 |
| 172 | CTTCTCCAGGTT | 116 |
| 173 | AGGACGGTTCTC | 117 |
| 174 | TATTCTGCCTCA | 118 |
| 175 | TCTGGTCTGCTT | 119 |
| 176 | GGAAGTTGTGAA | 120 |
| 177 | ACCTTGTATTCC | 121 |
| 179 | TGGAACTGTGTT | 122 |
| 180 | CAAGAGAAGGCC | 123 |
| 181 | CTTGTTCATTGC | 124 |
| 182 | CACACCTCTGCA | 125 |
| 188 | GGCACAACAGAA | 126 |
| 189 | AGAACACCTCGA | 127 |
| 190 | AATTAAGCTCCG | 128 |
| 192 | AGGATGAGTGTG | 129 |
| 196 | AGGTTATGGTGA | 130 |
| 197 | AGTATTCTCTGG | 131 |
| 198 | TATCCATTGTGG | 132 |
| 199 | CTTGTTCCACAA | 133 |
| 200 | CGAGCGGTGTTA | 134 |

**Supplementary Table 2:** Final 134 barcode sequences established as suitable for *in vivo* mRNA LNP screening. Listed index numbers correspond to LNP formulation numbers.

| Index | Ionizable lipid | %mol | Helper lipid | %mol | Cholesterol (%mol) | PEG-lipid | %mol |
|---|---|---|---|---|---|---|---|
| 1 | 1D4 | 35 | DOPE | 16 | 46.5 | C14-PEG2000 | 2.5 |
| 2 | 1D6.2 | 35 | DOPE | 16 | 46.5 | C14-PEG2000 | 2.5 |
| 3 | 1D8 | 35 | DOPE | 16 | 46.5 | C14-PEG2000 | 2.5 |
| 4 | 1D8i | 35 | DOPE | 16 | 46.5 | C14-PEG2000 | 2.5 |
| 5 | 1D18 | 35 | DOPE | 16 | 46.5 | C14-PEG2000 | 2.5 |
| 6 | 1D9.2 | 35 | DOPE | 16 | 46.5 | C14-PEG2000 | 2.5 |
| 7 | 2D4 | 35 | DOPE | 16 | 46.5 | C14-PEG2000 | 2.5 |
| 8 | 2D6.2 | 35 | DOPE | 16 | 46.5 | C14-PEG2000 | 2.5 |
| 9 | 2D8 | 35 | DOPE | 16 | 46.5 | C14-PEG2000 | 2.5 |
| 10 | 2D8i | 35 | DOPE | 16 | 46.5 | C14-PEG2000 | 2.5 |
| 11 | 2D18 | 35 | DOPE | 16 | 46.5 | C14-PEG2000 | 2.5 |
| 12 | 2D9.2 | 35 | DOPE | 16 | 46.5 | C14-PEG2000 | 2.5 |
| 13 | 3D4 | 35 | DOPE | 16 | 46.5 | C14-PEG2000 | 2.5 |
| 14 | 3D6.2 | 35 | DOPE | 16 | 46.5 | C14-PEG2000 | 2.5 |
| 15 | 3D8 | 35 | DOPE | 16 | 46.5 | C14-PEG2000 | 2.5 |
| 16 | 3D8i | 35 | DOPE | 16 | 46.5 | C14-PEG2000 | 2.5 |
| 17 | 3D18 | 35 | DOPE | 16 | 46.5 | C14-PEG2000 | 2.5 |
| 18 | 3D9.2 | 35 | DOPE | 16 | 46.5 | C14-PEG2000 | 2.5 |
| 19 | 4D4 | 35 | DOPE | 16 | 46.5 | C14-PEG2000 | 2.5 |
| 20 | 4D6.2 | 35 | DOPE | 16 | 46.5 | C14-PEG2000 | 2.5 |
| 21 | 4D8 | 35 | DOPE | 16 | 46.5 | C14-PEG2000 | 2.5 |
| 22 | 4D8i | 35 | DOPE | 16 | 46.5 | C14-PEG2000 | 2.5 |
| 23 | 4D18 | 35 | DOPE | 16 | 46.5 | C14-PEG2000 | 2.5 |
| 24 | 4D9.2 | 35 | DOPE | 16 | 46.5 | C14-PEG2000 | 2.5 |
| 25 | 5D4 | 35 | DOPE | 16 | 46.5 | C14-PEG2000 | 2.5 |
| 26 | 5D6.2 | 35 | DOPE | 16 | 46.5 | C14-PEG2000 | 2.5 |
| 27 | 5D8 | 35 | DOPE | 16 | 46.5 | C14-PEG2000 | 2.5 |
| 28 | 5D8i | 35 | DOPE | 16 | 46.5 | C14-PEG2000 | 2.5 |
| 29 | 5D18 | 35 | DOPE | 16 | 46.5 | C14-PEG2000 | 2.5 |
| 30 | 5D9.2 | 35 | DOPE | 16 | 46.5 | C14-PEG2000 | 2.5 |
| 31 | 6D4 | 35 | DOPE | 16 | 46.5 | C14-PEG2000 | 2.5 |
| 32 | 6D6.2 | 35 | DOPE | 16 | 46.5 | C14-PEG2000 | 2.5 |
| 33 | 6D8 | 35 | DOPE | 16 | 46.5 | C14-PEG2000 | 2.5 |
| 34 | 6D8i | 35 | DOPE | 16 | 46.5 | C14-PEG2000 | 2.5 |
| 35 | 6D18 | 35 | DOPE | 16 | 46.5 | C14-PEG2000 | 2.5 |
| 36 | 6D9.2 | 35 | DOPE | 16 | 46.5 | C14-PEG2000 | 2.5 |
| 37 | 7D4 | 35 | DOPE | 16 | 46.5 | C14-PEG2000 | 2.5 |
| 38 | 7D6.2 | 35 | DOPE | 16 | 46.5 | C14-PEG2000 | 2.5 |
| 39 | 7D8 | 35 | DOPE | 16 | 46.5 | C14-PEG2000 | 2.5 |
| 40 | 7D8i | 35 | DOPE | 16 | 46.5 | C14-PEG2000 | 2.5 |
| 41 | 7D18 | 35 | DOPE | 16 | 46.5 | C14-PEG2000 | 2.5 |
| 42 | 7D9.2 | 35 | DOPE | 16 | 46.5 | C14-PEG2000 | 2.5 |
| 43 | 8D4 | 35 | DOPE | 16 | 46.5 | C14-PEG2000 | 2.5 |
| 44 | 8D6.2 | 35 | DOPE | 16 | 46.5 | C14-PEG2000 | 2.5 |
| 45 | 8D8 | 35 | DOPE | 16 | 46.5 | C14-PEG2000 | 2.5 |
| 46 | 8D8i | 35 | DOPE | 16 | 46.5 | C14-PEG2000 | 2.5 |
| 47 | 8D18 | 35 | DOPE | 16 | 46.5 | C14-PEG2000 | 2.5 |
| 48 | 8D9.2 | 35 | DOPE | 16 | 46.5 | C14-PEG2000 | 2.5 |
| 49 | 10D4 | 35 | DOPE | 16 | 46.5 | C14-PEG2000 | 2.5 |
| 50 | 10D6.2 | 35 | DOPE | 16 | 46.5 | C14-PEG2000 | 2.5 |
| 51 | 10D8 | 35 | DOPE | 16 | 46.5 | C14-PEG2000 | 2.5 |
| 52 | 10D8i | 35 | DOPE | 16 | 46.5 | C14-PEG2000 | 2.5 |
| 53 | 10D18 | 35 | DOPE | 16 | 46.5 | C14-PEG2000 | 2.5 |
| 54 | 10D9.2 | 35 | DOPE | 16 | 46.5 | C14-PEG2000 | 2.5 |
| 55 | 11D4 | 35 | DOPE | 16 | 46.5 | C14-PEG2000 | 2.5 |
| 56 | 11D6.2 | 35 | DOPE | 16 | 46.5 | C14-PEG2000 | 2.5 |

| Index | Ionizable lipid | %mol | Helper lipid | %mol | Cholesterol (%mol) | PEG-lipid | %mol |
|---|---|---|---|---|---|---|---|
| 57 | 11D8 | 35 | DOPE | 16 | 46.5 | C14-PEG2000 | 2.5 |
| 58 | 11D8i | 35 | DOPE | 16 | 46.5 | C14-PEG2000 | 2.5 |
| 59 | 11D18 | 35 | DOPE | 16 | 46.5 | C14-PEG2000 | 2.5 |
| 60 | 11D9.2 | 35 | DOPE | 16 | 46.5 | C14-PEG2000 | 2.5 |
| 61 | 13D4 | 35 | DOPE | 16 | 46.5 | C14-PEG2000 | 2.5 |
| 62 | 13D6.2 | 35 | DOPE | 16 | 46.5 | C14-PEG2000 | 2.5 |
| 63 | 13D8 | 35 | DOPE | 16 | 46.5 | C14-PEG2000 | 2.5 |
| 64 | 13D8i | 35 | DOPE | 16 | 46.5 | C14-PEG2000 | 2.5 |
| 65 | 13D18 | 35 | DOPE | 16 | 46.5 | C14-PEG2000 | 2.5 |
| 66 | 13D9.2 | 35 | DOPE | 16 | 46.5 | C14-PEG2000 | 2.5 |
| 67 | 12D4 | 35 | DOPE | 16 | 46.5 | C14-PEG2000 | 2.5 |
| 68 | 12D6.2 | 35 | DOPE | 16 | 46.5 | C14-PEG2000 | 2.5 |
| 69 | 12D8 | 35 | DOPE | 16 | 46.5 | C14-PEG2000 | 2.5 |
| 70 | 12D8i | 35 | DOPE | 16 | 46.5 | C14-PEG2000 | 2.5 |
| 71 | 12D18 | 35 | DOPE | 16 | 46.5 | C14-PEG2000 | 2.5 |
| 72 | 12D9.2 | 35 | DOPE | 16 | 46.5 | C14-PEG2000 | 2.5 |
| 73 | 14D4 | 35 | DOPE | 16 | 46.5 | C14-PEG2000 | 2.5 |
| 74 | 14D6.2 | 35 | DOPE | 16 | 46.5 | C14-PEG2000 | 2.5 |
| 75 | 14D8 | 35 | DOPE | 16 | 46.5 | C14-PEG2000 | 2.5 |
| 76 | 14D8i | 35 | DOPE | 16 | 46.5 | C14-PEG2000 | 2.5 |
| 77 | 14D18 | 35 | DOPE | 16 | 46.5 | C14-PEG2000 | 2.5 |
| 78 | 14D9.2 | 35 | DOPE | 16 | 46.5 | C14-PEG2000 | 2.5 |
| 79 | 15D4 | 35 | DOPE | 16 | 46.5 | C14-PEG2000 | 2.5 |
| 80 | 15D6.2 | 35 | DOPE | 16 | 46.5 | C14-PEG2000 | 2.5 |
| 81 | 15D8 | 35 | DOPE | 16 | 46.5 | C14-PEG2000 | 2.5 |
| 82 | 15D8i | 35 | DOPE | 16 | 46.5 | C14-PEG2000 | 2.5 |
| 83 | 15D18 | 35 | DOPE | 16 | 46.5 | C14-PEG2000 | 2.5 |
| 84 | 15D9.2 | 35 | DOPE | 16 | 46.5 | C14-PEG2000 | 2.5 |
| 85 | 16D4 | 35 | DOPE | 16 | 46.5 | C14-PEG2000 | 2.5 |
| 86 | 16D6.2 | 35 | DOPE | 16 | 46.5 | C14-PEG2000 | 2.5 |
| 87 | 16D8 | 35 | DOPE | 16 | 46.5 | C14-PEG2000 | 2.5 |
| 88 | 16D8i | 35 | DOPE | 16 | 46.5 | C14-PEG2000 | 2.5 |
| 89 | 16D18 | 35 | DOPE | 16 | 46.5 | C14-PEG2000 | 2.5 |
| 90 | 16D9.2 | 35 | DOPE | 16 | 46.5 | C14-PEG2000 | 2.5 |
| 91 | 17D4 | 35 | DOPE | 16 | 46.5 | C14-PEG2000 | 2.5 |
| 92 | 17D6.2 | 35 | DOPE | 16 | 46.5 | C14-PEG2000 | 2.5 |
| 93 | 17D8 | 35 | DOPE | 16 | 46.5 | C14-PEG2000 | 2.5 |
| 94 | 17D8i | 35 | DOPE | 16 | 46.5 | C14-PEG2000 | 2.5 |
| 95 | 17D18 | 35 | DOPE | 16 | 46.5 | C14-PEG2000 | 2.5 |
| 96 | 17D9.2 | 35 | DOPE | 16 | 46.5 | C14-PEG2000 | 2.5 |
| 97 | 18D4 | 35 | DOPE | 16 | 46.5 | C14-PEG2000 | 2.5 |
| 98 | 18D6.2 | 35 | DOPE | 16 | 46.5 | C14-PEG2000 | 2.5 |
| 99 | 18D8 | 35 | DOPE | 16 | 46.5 | C14-PEG2000 | 2.5 |
| 100 | 18D8i | 35 | DOPE | 16 | 46.5 | C14-PEG2000 | 2.5 |
| 101 | 18D18 | 35 | DOPE | 16 | 46.5 | C14-PEG2000 | 2.5 |
| 102 | 18D9.2 | 35 | DOPE | 16 | 46.5 | C14-PEG2000 | 2.5 |
| 103 | 19D4 | 35 | DOPE | 16 | 46.5 | C14-PEG2000 | 2.5 |
| 104 | 19D6.2 | 35 | DOPE | 16 | 46.5 | C14-PEG2000 | 2.5 |
| 105 | 19D8 | 35 | DOPE | 16 | 46.5 | C14-PEG2000 | 2.5 |
| 106 | 19D8i | 35 | DOPE | 16 | 46.5 | C14-PEG2000 | 2.5 |
| 107 | 19D18 | 35 | DOPE | 16 | 46.5 | C14-PEG2000 | 2.5 |
| 108 | 19D9.2 | 35 | DOPE | 16 | 46.5 | C14-PEG2000 | 2.5 |
| 109 | 20D4 | 35 | DOPE | 16 | 46.5 | C14-PEG2000 | 2.5 |
| 110 | 20D6.2 | 35 | DOPE | 16 | 46.5 | C14-PEG2000 | 2.5 |
| 111 | 20D8 | 35 | DOPE | 16 | 46.5 | C14-PEG2000 | 2.5 |
| 112 | 20D8i | 35 | DOPE | 16 | 46.5 | C14-PEG2000 | 2.5 |

| Index | Ionizable lipid | %mol | Helper lipid | %mol | Cholesterol (%mol) | PEG-lipid | %mol |
|---|---|---|---|---|---|---|---|
| 113 | 20D18 | 35 | DOPE | 16 | 46.5 | C14-PEG2000 | 2.5 |
| 114 | 20D9.2 | 35 | DOPE | 16 | 46.5 | C14-PEG2000 | 2.5 |
| 115 | 9D4 | 35 | DOPE | 16 | 46.5 | C14-PEG2000 | 2.5 |
| 116 | 9D6.2 | 35 | DOPE | 16 | 46.5 | C14-PEG2000 | 2.5 |
| 117 | 9D8 | 35 | DOPE | 16 | 46.5 | C14-PEG2000 | 2.5 |
| 118 | 9D8i | 35 | DOPE | 16 | 46.5 | C14-PEG2000 | 2.5 |
| 119 | 9D18 | 35 | DOPE | 16 | 46.5 | C14-PEG2000 | 2.5 |
| 120 | 9D9.2 | 35 | DOPE | 16 | 46.5 | C14-PEG2000 | 2.5 |
| 121 | C12-200 | 35 | DOPE | 16 | 46.5 | C14-PEG2000 | 2.5 |
| 122 | cKK-E12 | 35 | DOPE | 16 | 46.5 | C14-PEG2000 | 2.5 |
| 123 | DLin-MC3-DMA | 50 | DSPC | 10 | 38.5 | DMG-PEG2000 | 1.5 |
| 124 | 306Oi10 | 35 | DOPE | 16 | 46.5 | C14-PEG2000 | 2.5 |
| 125 | SM-102 | 50 | DSPC | 10 | 38.5 | DMG-PEG2000 | 1.5 |
| 126 | ALC-0315 | 46.3 | DSPC | 9.4 | 42.7 | ALC-0159 | 1.6 |
| 127 | C14-482 | 35 | DOPE | 16 | 46.5 | C14-PEG2000 | 2.5 |
| 128 | C14-488 | 35 | DOPE | 16 | 46.5 | C14-PEG2000 | 2.5 |
| 129 | C16-488 | 35 | DOPE | 16 | 46.5 | C14-PEG2000 | 2.5 |
| 130 | C12-494 | 35 | DOPE | 16 | 46.5 | C14-PEG2000 | 2.5 |
| 131 | C14-494 | 35 | DOPE | 16 | 46.5 | C14-PEG2000 | 2.5 |
| 132 | C16-494 | 35 | DOPE | 16 | 46.5 | C14-PEG2000 | 2.5 |
| 133 | C14-c494 | 35 | DOPE | 16 | 46.5 | C14-PEG2000 | 2.5 |

**Supplementary Table 3:** Formulation details of tested LNP library.

| Marker | Clone |
|---|---|
| CD3 | 17A2 |
| CD19 | 6D5 |
| CD11b | M1/70 |
| CD11c | N418 |
| CD31 | MEC13.3 |
| CD45 | 30-F11 |
| CD68 | FA-11 |
| CD146 | ME-9F1 |
| CD326 | G8.8 |
| CD335 | 29A1.4 |
| F4/80 | BM8 |

**Supplementary Table 4:** Antibody clones used for flow cytometry and FACS.

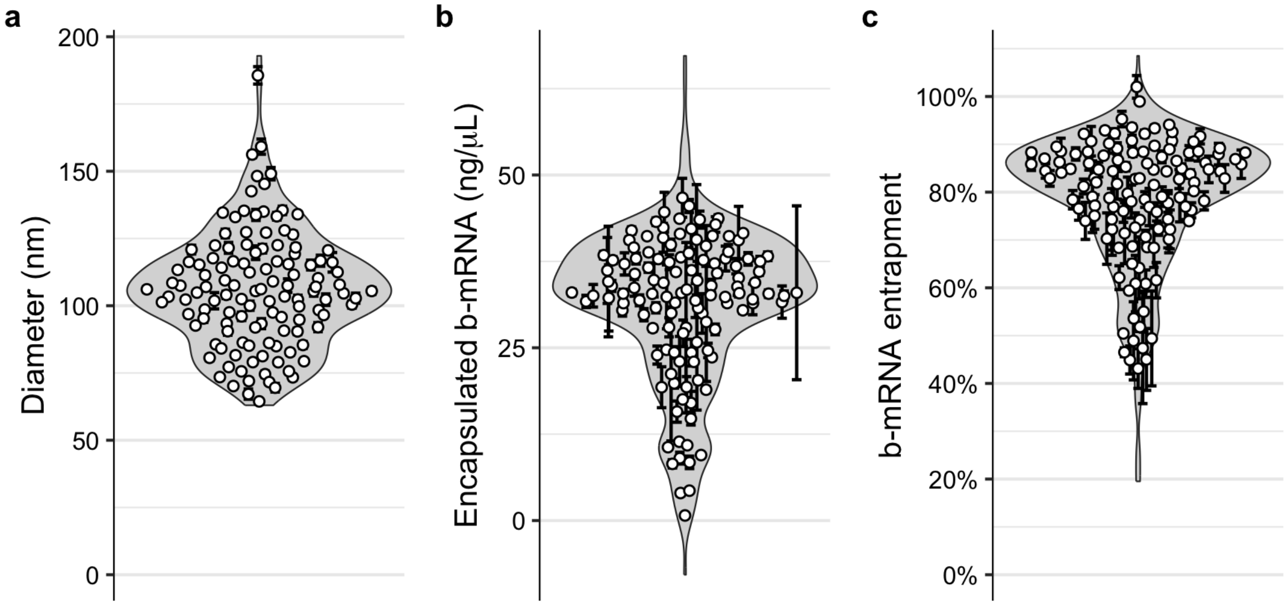

**Supplementary Figure 1:** Physicochemical characterization of b-mRNA LNP library. **a.** Hydrodynamic LNP diameter as measured by DLS. **b.** Encapsulated b-mRNA concentration as measured by RiboGreen. **c.** b-mRNA entrapment efficiency as measured by RiboGreen. Data are presented as mean ± standard error of the mean from $n \geq 4$ independent measurements.

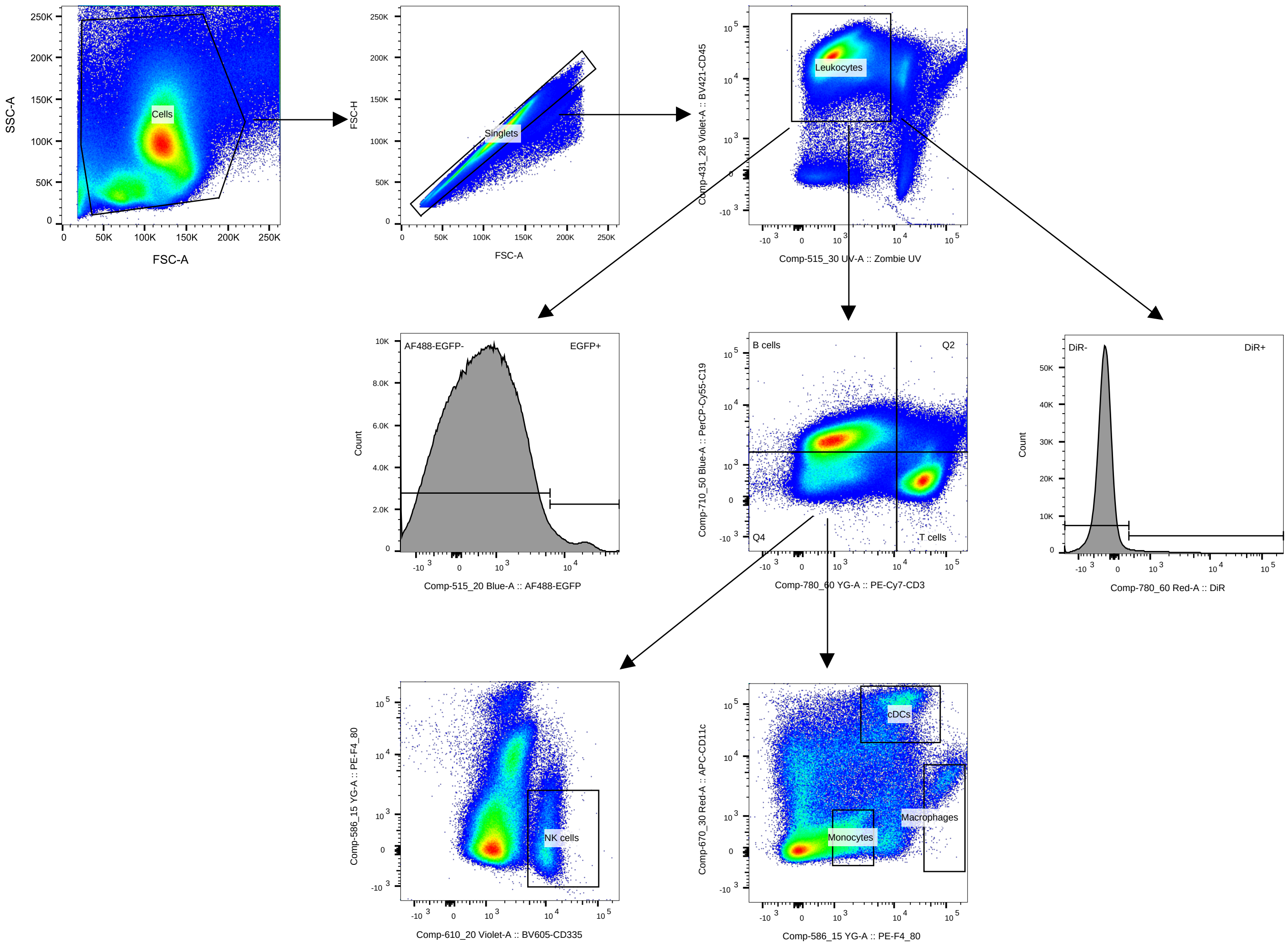

**Supplementary Figure 2:** Representative blood/spleen flow cytometry gating scheme used for initial FACS-based b-mRNA LNP screen and for subsequent validation experiments.

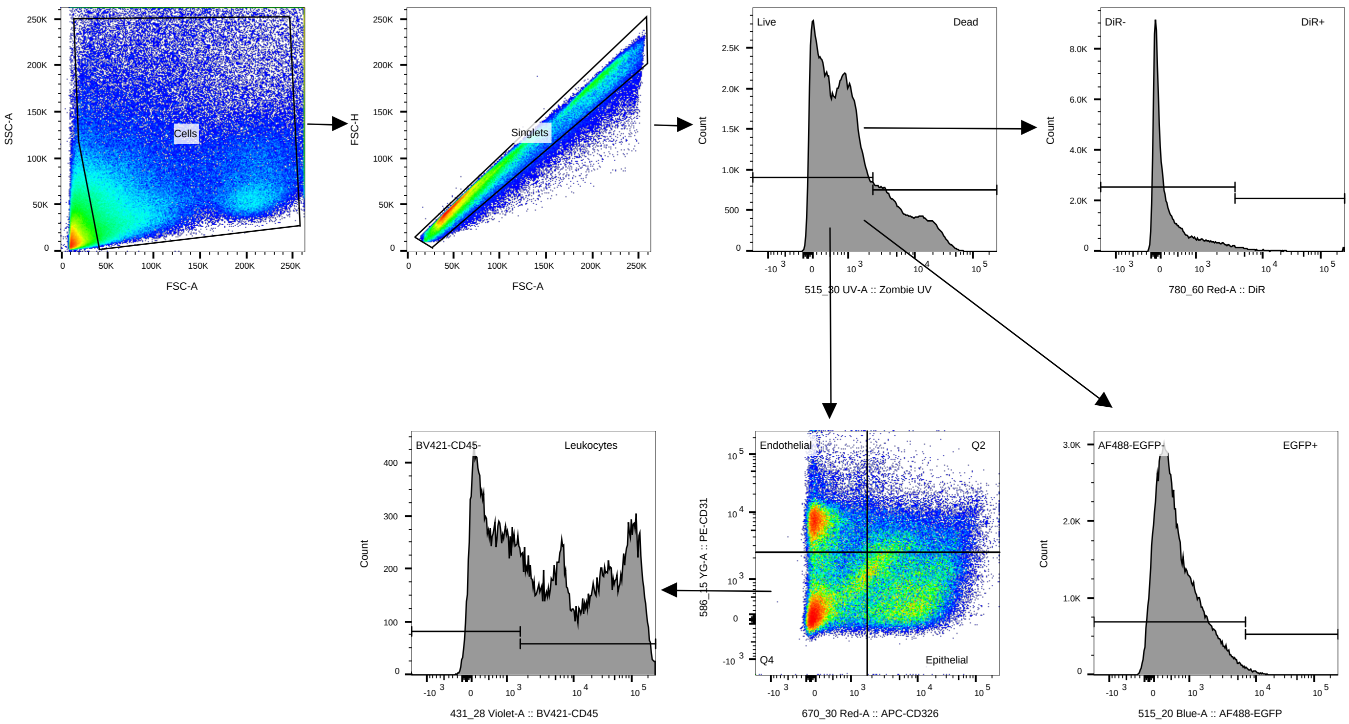

**Supplementary Figure 3:** Representative lung flow cytometry gating scheme used for initial FACS-based b-mRNA LNP screen and for subsequent validation experiments.

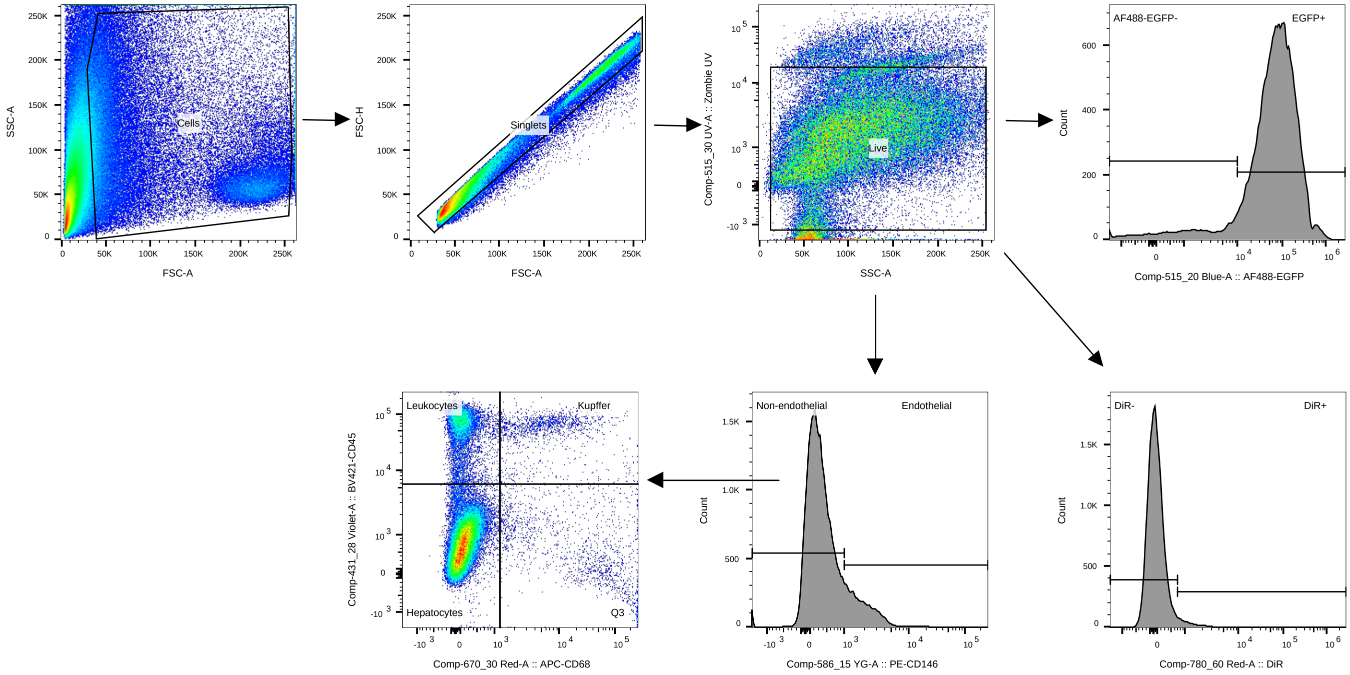

**Supplementary Figure 4:** Representative liver flow cytometry gating scheme used for initial FACS-based b-mRNA LNP screen and for subsequent validation experiments.

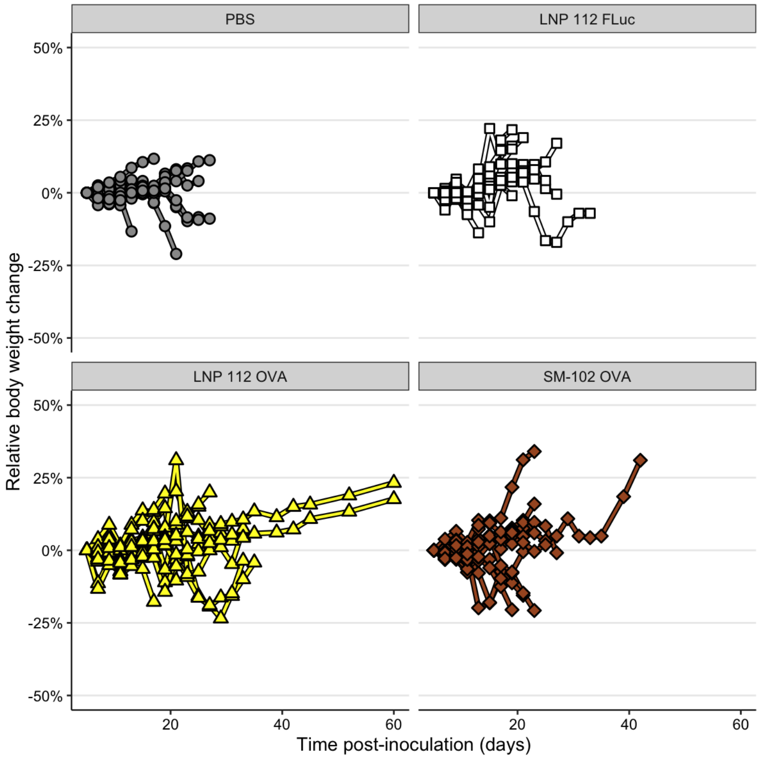

**Supplementary Figure 5:** Body weight change of mice inoculated with B16-OVA melanoma cells and given the indicated treatments in a therapeutic cancer vaccine model.
